# IL-1β/IRAK4 Axis Promotes Ovarian Tumor Development at the Mesothelium Injury Sites

**DOI:** 10.1101/2025.03.18.643950

**Authors:** John P. Miller, Kyu Kwang Kim, Negar Khazan, Cameron WA Snyder, Niloy A. Singh, Elizabeth Lamere, Myla Strawderman, Sonali Sharma, Ronald Lakony, Michelle Whittum, Mark Anderson, Rick Keenan, Elizabeth Pritchett, Cameron Baker, John Ashton, Manoj K. Khera, Charles C. Chu, Jeevisha Bajaj, Michael R Elliott, Rachael Rowswell-Turner, Laura M Calvi, Michael W. Becker, Richard G. Moore, Rakesh K. Singh

## Abstract

Epithelial ovarian cancer (EOC) cells seed at mesothelial inflammation or injury sites. Lack of animal models recapitulating tumor cells seeding at inflamed sites in EOC hinders mechanistic studies and therapy developments. Here, we developed a non-surgical MIM (Mesothelium Inflammation/Injury Metastasis) model that recapitulates tumor cell seeding at inflamed sites. This model captures temporal changes in tumor immune microenvironment and tumor growth allowing for deeper mechanistic and preclinical therapeutic studies of EOC *in-vivo*. We show here that HGS-3 high-grade murine serous EOC cells seed at needle-induced injury sites in mesothelium/peritoneal wall, forming tumors both internally and protruding outward. Using MIM model, we found that deletion of IL1R1 in mice reduced EOC cell seeding at mesothelium injury/inflamed site in WT but not IL1ra-deficient mice. Treatment of MiM mice with a novel IRAK4 inhibitor we recently developed (UR241-2) revealed an essential role for IRAK4 signaling downstream IL-1β/IL-1R1 in fostering an anti-tumor inflammatory environment, and reduced tumor burden. We conclude that IRAK4 inhibitors can be more effective than IL-1/IL-1R1 targeting agents to control metastasis and peritoneal tumors, an unmet medical need in EOC recurrence. Downregulation of extracellular matrix (ECM), upregulation of neutrophil activation genes, reduced cell adhesion and migration exhibit how UR241-2 corrects ECM and immune disorders in EOC, making it less conducive to metastasis and tumorigenesis.

## Introduction

Inflammation can support the development of epithelial ovarian cancer (EOC)^1–2^ and is shown to be involved in the progression of various other malignancies^3–4^. Inflammation recruits the IL1β/TLR/IRAK4 axis, activating the downstream transcription factor NF-κβ ^5–7^. In EOC, IL1β^8–10^, its receptor IL1R^11–12^, the myddosome complex partners TLR4/Myd88^13–18^, IRAK4^19–20^, and NF-κβ^21–30^ are each aberrantly overexpressed, predicting poor survival and chemoresistance^31^. Notably, IL1β plays a critical role in peritoneal dissemination^32^, loss of mesothelial integrity^33–34^, conversion of ascitic tumor cells into peritoneal solid tumors, and the enrichment of malignant cytokines in peritoneal effusions^35–39^. These processes directly implicate IL1β in tumor cell adhesion and metastasis—the primary cause of EOC lethality. However, therapeutic strategies to block IL1β-driven metastatic events have yet to emerge for malignancies, including EOC. Despite IL1β’s compelling role in EOC cell seeding, Anakinra, a recombinant IL-1R1 antagonist that competitively inhibits IL-1β binding to IL-1R1, and Canakinumab^40^, an IL-1β-targeting antibody, have not been evaluated as potential therapies to prevent metastasis. There is a critical unmet medical need for strategies to block metastasis, particularly in recurrent disease, which is characterized by widespread metastases, chemoresistance, and high mortality in EOC. Additionally, the roles of IRAK1 and IRAK4—key downstream kinases mediating IL-1β/TLR/IRAK4 signaling^41–43^—remain unexplored in the context of IL-1β-driven cell seeding and ovarian tumorigenesis. Targeted therapies against both IRAK1 and IRAK4 for EOC treatment are also lacking.

In this study, we investigated the roles of IL1β/IL-1R1, IRAK1/4, and NLRP3 in cell adhesion at peritoneal mesothelial injury sites and the omentum (non-injury site). Using our MiM model, we assessed the impact of IL1R1, its antagonist IL1RN, and NLRP3 on seeding outcomes. To do this, we employed *Il-1r1*^KO^, *Il-1ra*^KO^, and *Nlrp3*^KO^ mice, intraperitoneally implanted with HGS-3 murine high-grade serous EOC cells, which recapitulate the fallopian tube origin of high-grade serous EOC. In addition, we primarily focused on elucidating the roles of IRAK4 in seeding using the MIM model and on developing a novel, selective IRAK4 small-molecule inhibitor, UR241-2. We fully characterized the pharmacology and absorption, distribution, metabolism and excretion (ADME) of UR241-2 for clinical translation and examined the impact of IRAK4 on: (1) extracellular matrix (ECM) elements, (2) the tumor immune milieu, (3) global genomic changes via bulk RNA-seq of vehicle- and UR241-2-treated tumors to identify affected genes/pathways, and (4) overall ovarian tumor growth using SKOV-3 EOC xenografts and a syngeneic murine high-grade serous EOC model that recapitulates the fallopian tube origin of EOC.

Taken together, our study uncovered key mechanisms driving EOC cell seeding at inflamed or injured mesothelial sites and identified pathways that, when targeted with IRAK4 inhibitors could control metastasis alone or potentially greater upon combining with inhibitors of the proteins and pathways involved in metastasis. By probing the role of IRAK4 in adhesion and metastasis, we provide insights that support IRAK4 inhibitors as potential treatments, particularly in recurrent EOC, where unmet medical needs are greatest.

## Results

### Il1β/Il1r1^LOF^ reduces seeding of murine high-grade serous EOC at the needle injury site, the development of murine metastatic MiM model

Analysis of EOC patient microarray data revealed that elevated IL-1β and IL-1R1 expression predicts poor prognosis (Fig-1A-B). To investigate the roles of IL-1β/IL-1R1 in ovarian tumorigenesis, particularly in metastasis—a process driven by IL-1β—we established a metastatic murine MIM model (See Methods Section). HGS-3 murine high-grade EOC cells (3.7 million/mice) were injected intraperitoneally (IP) into C57BL/6 mice using 21-gauge needles. By 30–45 days post-inoculation, tumor nodules emerged at the site of needle injury. To assess the role of IL-1R1, we implanted a similar number of HGS-3 cells in both C57BL/6 wild-type (WT) and *Il1r1* knockout (*Il1r1*^KO^; B6.129S7-Il1r1tm1Imx/J) mice (Fig-1C). After 42 days, mice were euthanized, and tumors at the injury site and omentum were harvested and weighed. WT mice developed cutaneous tumor nodules in 80% of cases (Fig-1D). In contrast, *Il1r1*^KO^ mice exhibited reduced tumor incidence and smaller cutaneous tumors (Fig-1E), while omental tumor frequency and weight remained comparable between WT and *Il1r1*^KO^ mice (Fig-1F). Notably, tumor weights at inflamed cutaneous sites differed significantly (Fig-1G). To confirm that *Il1r1*^KO^ reduces peritoneal tumors (Fig-1E), we implanted HGS-3 cells in *Il1rn* knockout (*Il1rn*^KO^) mice. WT and *Il1rn*^KO^ mice^44^ developed omental (Fig-1I) and skin tumors (Fig-1L) of similar size (Fig-1J, M). Next, we examined whether *Nlrp3*, a key inflammasome component, influences cell adhesion and tumor formation similarly to *Il1r1*. HGS-3 cells were implanted in *Nlrp3*^KO^ (B6.129S6-Nlrp3tm1Bhk/J) mice. Tumor burden at both cutaneous and omental sites in *Nlrp3*^KO^ mice mirrored that of WT mice (Supplementary Fig-1).

**Figure-1:**
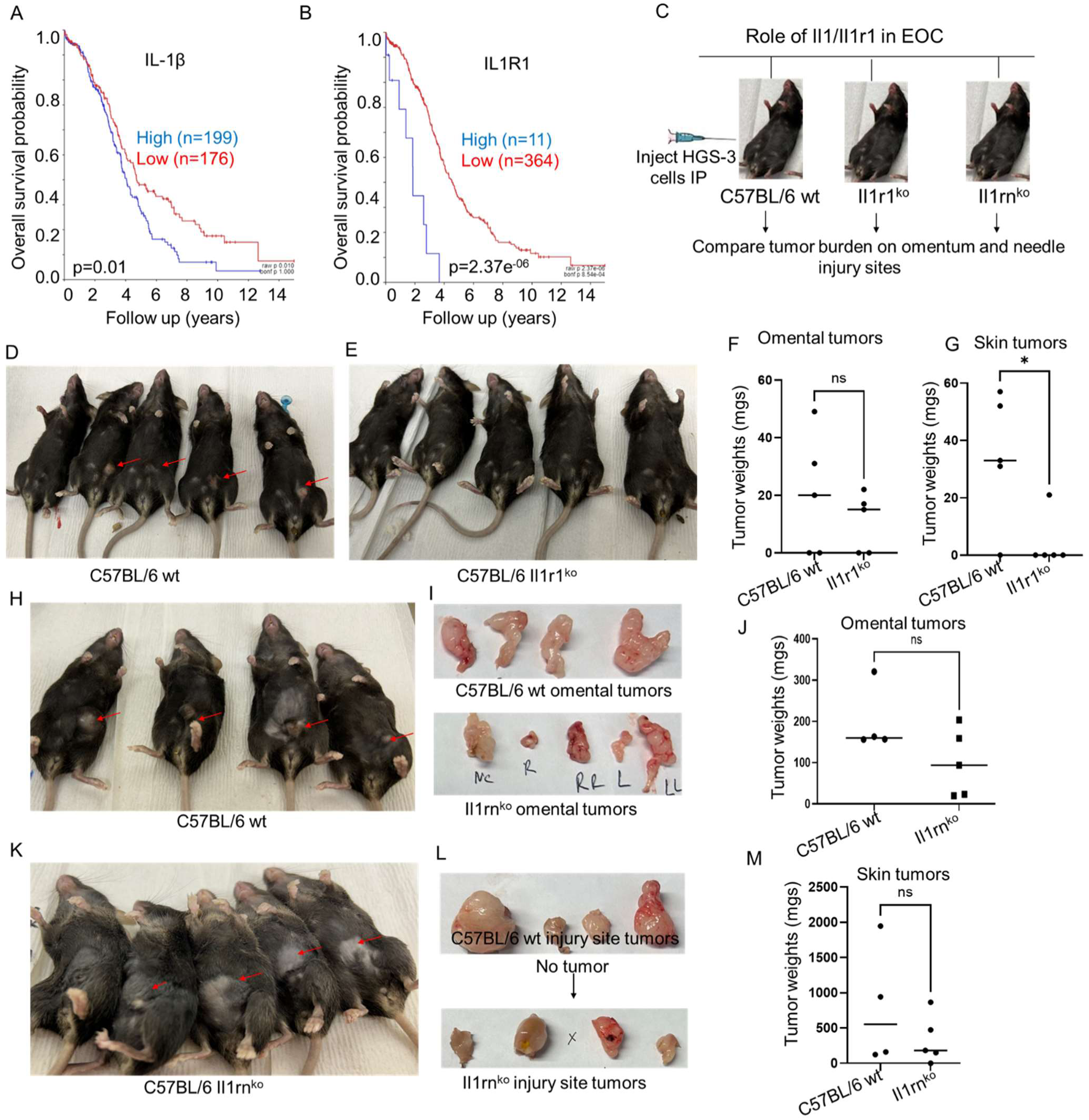
(**A-B**): Analysis of microarray data (TCGA-381-tpm-gencode36) of ovarian cancer patients using R2 Genomics Analysis and Visualization Platform tools showed that IL1β and IL1R1 mRNA overexpression predicts poor survival. (**C**): Schema of testing the role of IL1β, IL1R1 and IL1RN (IL1R1 antagonist) in ovarian cancer burden both at the needle injury and omental sites. (**D-E**): HGS-3 murine high-grade serous EOC cells (3 million/per mice) were implanted intraperitoneally using 21 sterile gauge needle in C57BL/6 WT and C57BL/6 IL1R1^KO^ mice. Mice were observed for 45-50 days and euthanized. Tumors formed on needle injury site both protruding at the skin and, in the peritoneum, and on the omentum, shown by red arrows, were isolated, weighed and frozen in liquid nitrogen. Lavages via washing with sterile PBS(5mL) were also collected. The studies were repeated thrice. A representative experiment is shown. (**F**): Weights of the omental did not differ between C57BL/6 WT and IL1R1^KO^ mice. (**G**): Weights of the peritoneal /skin tumors differed significantly between C57BL/6 wt and IL1R1^KO^ mice groups. * indicates <0.05. (**H-K**): Tumors formed at the needle injury site in C57BL/6 WT and IL1RN^KO^ mice are pointed with the red arrows. (**I**-upper vs **I**-lower): Images of the omental and peritoneal /skin tumors harvested from the euthanized mice are shown. (**J**): Omental tumor weights in C57BL/6 wt did not differ between from IL1RNko mice groups. (**K**): Weights of the tumors formed on the needle injury site in C57BL/6 WT mice did not differ for IL1RN^KO^ mice groups. This experiment was repeated twice. Tumor sizes were analyzed via non-parametric T-test using Graph-Prism version -7 or higher. * indicates <0.05.

### IRAK1/4 associates with mortalities and EOC disease stages and invasion

Schema of IL-1R/TLR/IRAK1/4 signaling is shown (Fig-2A). Analysis of the EOC patient’s microarray of the data available with R2-Genomcis Analysis and Data Visualization platform showed that IRAK1 (p=3.21e^-03^), IRAK2 (p=0.062) and IRAK4 (p=4.8e^-06^), exhibit poor prognoses in EOC patients (Fig-2, B-D). Analysis of the EOC patient’s microarray of the data available with GENT2^45^ tools showed that compared to normal ovaries, IRAK4 mRNA was overexpressed in malignant ovaries (p<0.001, log_2_Fold(F):0.979) (Fig-2E). Analysis of the EOC patient’s microarray data using GENT2 tools showed that IRAK4 mRNA expression was altered depending on the stage of EOC. Two-Sample T-test showed the following changes: IA vs I (p=0.004); IA vs II (p=0.007); IA vs III (p<0.001); IC vs IIA (p=0.002); IC vs III (p<0.001); IIA vs I (p<0.001); IIA vs II (p<0.001); IIA vs III (p<0.001); IIA vs IIC (p=0.006); IIA vs IIIC (p=0.001); IIC vs III (p<0.001); III vs IIB (p=0.004); IIIA vs I (p=0.001); IIIA vs II (p=0.004); IIIA vs III (p<0.001); IIB vs I (p=0.008); IIIB vs III (p=<0.001); IV vs I (p=0.004); IV vs IIA (p=0.005); IV vs III (p=<0.001). (http://gent2.appex.kr/gent2/, date accessed 10/2/2023) (Fig-2F). Notably, compared to EOC patients presenting with no-invasion, those with invasion showed increased IRAK4 mRNA expression (p=4.92e^-03^) (Fig-2G).

**Figure-2:**
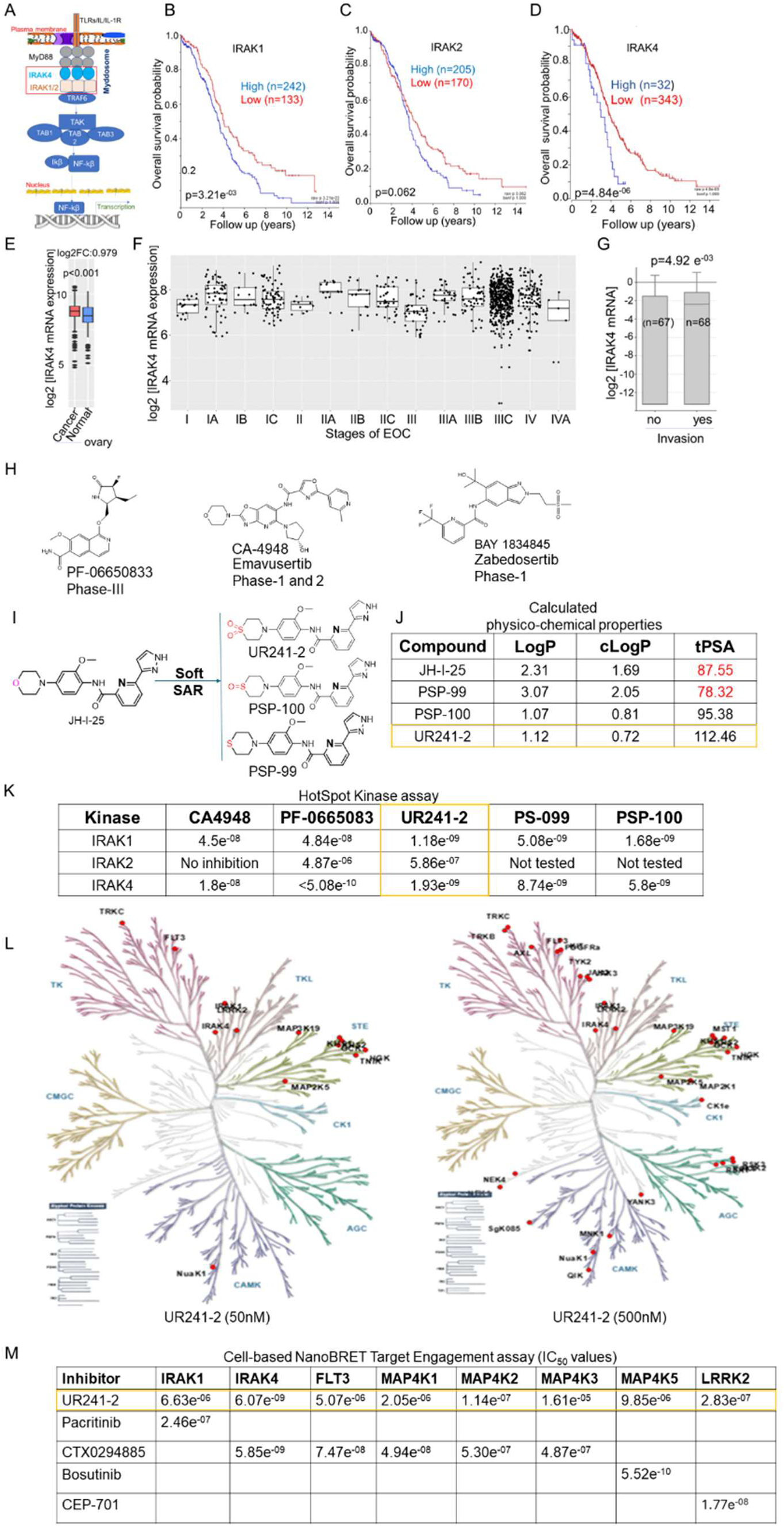
(**A**): Schema of IL1β/TLR/IRAK1/2/4 signaling pathway. (**B-D**): Analysis of ovarian cancer patient’s microarray data (TCGA-381-tpm-gencode36) using R2 Genomics Analysis and Visualization Platform tools showed that IRAK1, -2 and -4 mRNA overexpression predict poor survival. (**E**): IRAK4 mRNA was overexpressed in malignant than normal ovaries. The data present in GENT2 ovarian cancer databases were analyzed using the inbuilt tools. (**F**): Analysis of the EOC patient’s microarray data using GENT2 tools showed that IRAK4 mRNA expression was altered in various stages of disease. Two-sample T-test showed statistical differentiations: IA vs I (p=0.004); IA vs II (p=0.007); IA vs III (p<0.001); IC vs IIA (p=0.002); IC vs III (p<0.001); IIA vs I (p<0.001); IIA vs II (p<0.001); IIA vs III (p<0.001); IIA vs IIC (p=0.006); IIA vs IIIC (p=0.001); IIC vs III (p<0.001); III vs IIB (p=0.004); IIIA vs I (p=0.001); IIIA vs II (p=0.004); IIIA vs III (p<0.001); IIB vs I (p=0.008); IIIB vs III (p=<0.001); IV vs I (p=0.004); IV vs IIA (p=0.005); IV vs III (p=<0.001). (http://gent2.appex.kr/gent2/, date accessed 10/2/2023). (**G**): Compared to non-invasive EOC, tissues from invaded EOC phenotype showed greater IRAK4 mRNA. EOC patient microarray data (TCGA-541-custom-tcgaovag1) deposited at R2-Genomics Analysis and Visualization Platform were analyzed using their MegaSnitch tools. (**H**): Chemical structures of three representative IRAK4–kinase inhibitors undergoing clinical trials. (**I**): Structure-activity relationship (SAR) guided optimization of JH-I-25 scaffold, a literature described IRAK4 inhibitor leading to 3 potent novel analogs. The chemical structure of UR241-2 (sulfone), PSP-099 (sulfide) and PSP-100 (sulfoxide), from among the many analogs synthesized, are shown. (**J**): Calculated drug-likeness/physicochemical properties (LogP, cLogP and topological polar surface area-tPSA are shown. These properties were calculated using Chemdraw software. (**K**): Comparison of IRAK1-, IRAK2-, and IRAK4-kinase inhibitory IC50s of UR241-2 versus CA4948, PF-06650833, PSP-099 and PSP-100 are shown. Compounds were screened using HotSpot Kinase assay available in Reaction Biology Laboratories using 1μM ATP concentration under a 10-dose singlet screening program. (**L**): Dendrograms of global kinome activity of UR241-2 at 50- and 500nM doses. Red indicates kinase affected. At 50nM 13 of 682 kinases were significantly inhibited, whereas at 500nM 34 kinases, shown as red dots, from among 682 total kinases were inhibited. (**M**). A HEK293 cell-based NanoBret target Engagement screening assay was conducted to determine the selectivity of UR241-2 among 10 most affected kinases revealed by HotSpot kinase assay. The nanoBret assay showed that UR241-2 inhibits IRAK4 kinase activity only and other kinases including IRAK1 are affected at 10-100 folds higher doses, making UR241-2 as one of the most selective and specific IRAK4 kinase inhibitors known currently.

### UR241-2 is a highly selective and potent IRAK4 kinase inhibitor with minimal off-target activity

Several IRAK4 kinase inhibitors are in clinical development for various diseases, including cancer (Fig-2H). CA4948, a dual IRAK4/FLT3 inhibitor, is undergoing clinical trials for AML, pancreatic, esophageal, and gastric cancers. Given the association of EOC with IL-1/TLR/MYD88/IRAK4 signaling, targeting IRAK4 kinase presents a compelling therapeutic strategy. However, CA4948 also inhibits CLK, and since FLT3 and CLK upregulation in EOC correlates with better prognosis (Supplementary Fig-2). For our studies, we developed UR241-2, an IRAK4-selective inhibitor devoid of off-target effects of FLT3. We considered bio-isosteric modification of JH-I-25, an IRAK4 inhibitor with ∼20nM potency (add citation). We replaced the morpholine side chain in JH-I-25 with more physicochemical-activity-privileged bio-isosteres such as thiomorpholine, thiomorpholine-S-oxide, or thiomorpholine-S-dioxide (Fig-2I) to generate a novel compound with significantly improved clogP and total polar surface area (tPSA) values in silico (Fig-2J). When synthesized and screened via Hotspot kinase profiling (Reaction Biology Inc., PA, USA), the resulting compounds—UR241-2, PS-099, and PS-100— exhibited ∼10- to 12-fold superior IRAK1/4 inhibition compared to JH-I-25 (Fig-2K). UR241-2 demonstrated the strongest IRAK1/4 inhibition and was selected as the lead compound. To assess off-target effects, we conducted global kinase screenings at 50nM and 500nM doses. Dendrogram analysis revealed that only 13 kinases were affected at 50nM, and 34 kinases at 500nM (Fig-2L). Importantly, NanoBRET Intracellular Kinase Assay in intact HEK293 cells confirmed that IRAK4 was the only significantly affected target, underscoring UR241-2’s notable specificity and selectivity (Fig-2M)

### UR241-2 exhibits greater binding affinity and distinct interactions compared to JH-I-25

To elucidate the binding mode of UR241-2 and its analogs with IRAK4 and facilitate structure–activity relationship (SAR) development, we performed in silico docking simulations using UR241-2, PS-099, and PS-100 against the known IRAK4 crystal structure (Fig-3A). The interacting residues and nature of these interactions for each ligand are shown in Fig-3B, with JH-I-25 serving as a comparator. In these analyses, we examined aliphatic (green), aromatic (magenta), basic (blue), polar (sky blue), sulfur (yellow), hydrogen bond donor/acceptor (HBD/HBA) (dotted sky blue), hydrophobic (dotted green), and Van der Waals (VdW) (dotted cyan) interactions (Fig-3B, bottom). As expected from JH-I-25’s docking profile, the pyrazolo-pyridine core of all ligands occupied the same hinge region, though with modified residual interactions, highlighting a potential avenue for enhancing IRAK4 inhibition. Further analysis revealed that replacing the oxygen in JH-I-25’s morpholine with sulfur in thiomorpholine (PS-099) introduced a novel ASP114 hydrogen bond interaction, absent in JH-I-25. Although the pyrazole nitrogen atoms in JH-I-25 and PS-100 retained interactions with SER165, PHE34, ASN153, and ALA152, neither UR241-2 nor PS-099 engaged these residues, despite maintaining similar symmetry elements in their morpholine, thiomorpholine, sulfoxide, and sulfone groups (Fig-3B). A superimposed view of all four ligands docked together is shown in Fig-3C. Next, we conducted molecular dynamics simulations (MDS) to evaluate the dynamic behavior of each ligand within the IRAK4 catalytic site and validate the docking results (Fig-3D-H). MDS assessed key binding interactions through root-mean-square deviation (RMSD), root-mean-square fluctuation (RMSF), and radius of gyration, all measured in Å. RMSD quantifies ligand deviation from the crystal structure upon docking, while RMSD-protein measures structural deviation of IRAK4 itself. Our MDS studies demonstrated that all ligands maintained tight interactions within 2Å of IRAK4 when simulated over 1 nanosecond (ns), with an RMSD of <2Å considered highly stable (Fig-3D/E). RMSF, which represents fluctuations relative to the top pose, was also analyzed (Fig-3F). Energy calculations revealed the highest binding affinity for UR241-2 (sulfone), followed by PS-100 (sulfoxide) and JH-I-25, supporting our hypothesis. Interestingly, PS-099, the most non-polar analog, exhibited the lowest binding affinity (−10.5 kcal/mol) (Fig-3H/I).

**Figure-3:**
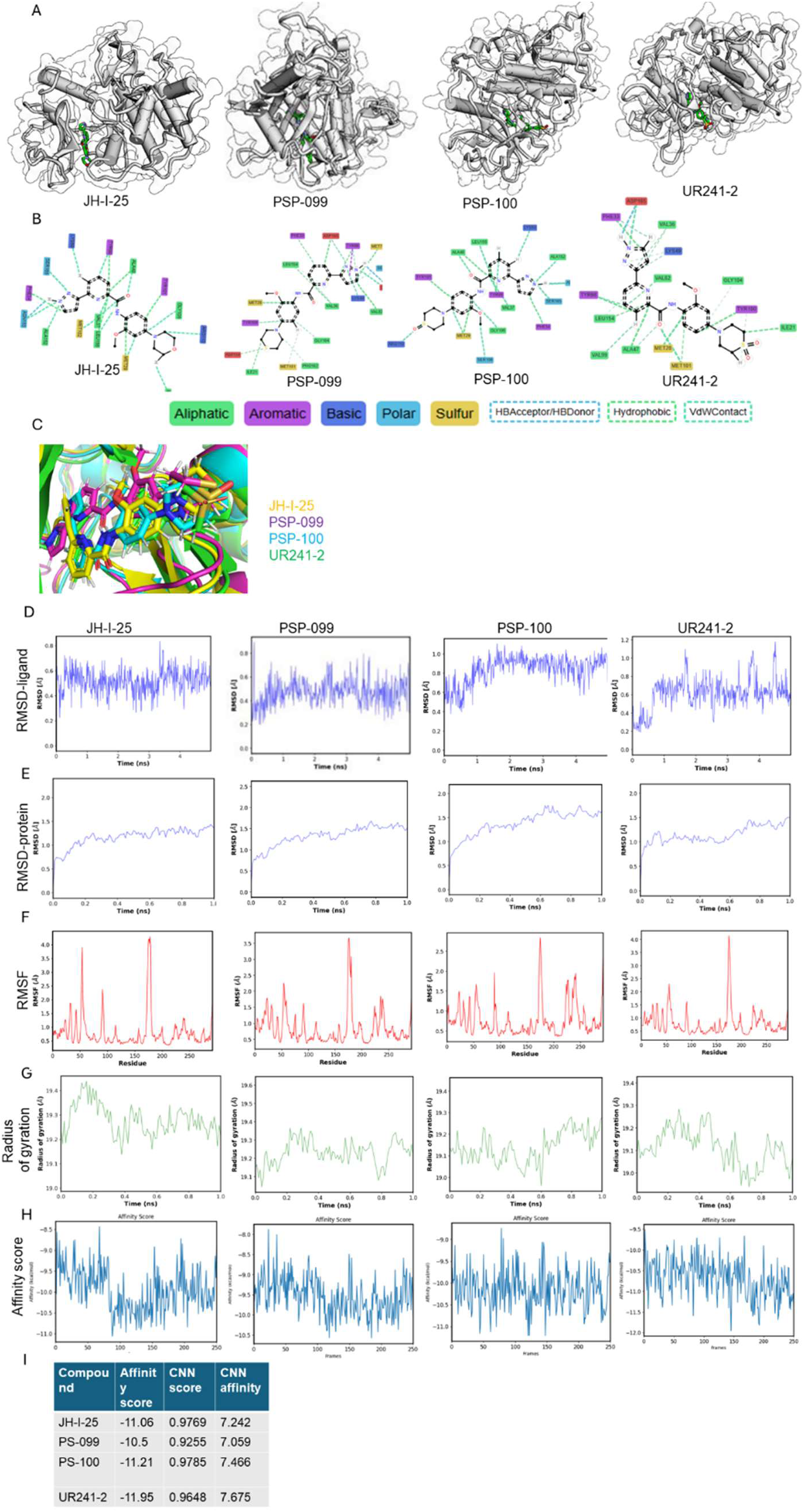
(**A**): JH-I-25 and its newer analogs UR241-2, PSP-099 and PSP-100 were docked individually to IRAK4 crystal structure using Gnina docking software, which is built on neural networks (CNNs) as a scoring function. (**B**): The interacting amino acid residues and their nature of interactions are shown. Green=aliphatic, magenta=aromatic, blue=basic, sky blue=polar, and yellow=sulfur. Red indicates unfavorable interactions. (**C**): JH-I-25, UR241-2, PSP-099 and PSP-100 analogs were docked together to IRAK4 protein using Gnina tools. (**D**): Root-mean square density (RMSD) simulations of JH-I-25, UR241-2, PSP-099 and PSP-100 ligands docked to IRAK4 falling within 2Å unit range are shown. RMSD-ligand is the root mean deviation of the bounded ligand compared to the ligand in the crystal structure. (**E**): Root-mean square density (RMSD) simulations of IRAK4 protein docked with JH-I-25, UR241-2, PSP-099 and PSP-100 ligands falling with 2Å unit range are shown. RMSD-Protein is the root mean deviation of the protein for each frame of the simulation compared to the crystal structure. Both **D** and **E** indicate that docking quality was acceptable. (**F**): RMSF (root mean square fluctuation) of JH-I-25, UR241-2, PSP-099 and PSP-100 ligands are shown. The ligands, when bound, induce the most notable structural changes in residues 172-181 of IRAK4 which reflected in the spike in the RMSF charts. RMSF is the mean fluctuation of IRAK4 protein residues after binding of ligands. RMSF indicates the conformational flexibility of the complex. The time/frame element is removed from RMSF in order to show an average fluctuation of the residues. (**H**): The affinity score measures affinity of a ligand (in that particular frame) binding to the protein. The frames represent different conformations that a ligand may adopt. The affinity is measured in KCal/mol and the lowest frame (lowest energy) is selected as the most stable binder. UR241-2 showed best affinity score (−11.95), hence was considered the most stable binder ligand of IRAK4 crystal structure.

### UR241-2 exhibits favorable ADMET (Absorption, Distribution, Metabolism, Excretion, and Toxicity) characteristics

We first evaluated kinetic solubility at pH 7.4, where UR241-2 demonstrated high solubility (29 µM) in phosphate-buffered saline (PBS) with 1% DMSO (Supplementary Fig-3A). Next, we assessed metabolic stability in human and mouse liver microsomes (HLM and MLM), a key determinant of hepatic clearance and bioavailability. UR241-2 exhibited high stability in HLM (t₁/₂ = 208.9 min, CLint = 3.33 µL/min/mg protein), whereas in MLM, it showed faster clearance (t₁/₂ = 8.70 min, CLint = 79.7 µL/min/mg protein) (Supplementary Fig-3B).To evaluate potential drug–drug interaction risks, we tested UR241-2 against eight cytochrome P450 (CYP) isoforms (1A2, 2B6, 2C8, 2C9, 2C19, 2D6, 3A4M, 3A4T). Weak inhibition was observed, with IC₅₀ values ranging from 30 to >100 µM, indicating minimal CYP liabilities (Supplementary Fig-3C). Plasma protein binding studies showed high binding efficiency in both human (97.4 ± 0.3%) and mouse (93.8 ± 0.8%) plasma (Supplementary Fig-3D). CaCo-2 permeability assays revealed a high efflux ratio (4.14), classifying UR241-2 as an efflux substrate (Supplementary Fig-3E). Finally, we assessed hERG (human Ether-à-go-go Related Gene) channel inhibition, a key predictor of cardiotoxicity. UR241-2 showed no significant hERG inhibition at doses up to 100 µM (EC₅₀ = 1.01 × 10⁻⁴), whereas the positive control E-4031 exhibited strong inhibition (EC₅₀ = 1.82 × 10⁻⁸) (Supplementary Fig-3F).

### UR241-2 demonstrates acceptable plasma levels in a mouse pharmacokinetic (PK) study

To assess serum levels over time, we conducted a pilot PK study in CD-1 mice. UR241-2 was administered at 20 mg/kg intraperitoneally (IP) and 1 mg/kg intravenously (IV). Plasma concentrations were analyzed using LC-MS. As shown in Supplementary Fig-3G, IV administration resulted in rapid clearance, whereas IP administration (20 mg/kg) maintained plasma levels above 8 ng/mL for up to 8 hours. This study provides critical data to guide dose selection for *in vivo* experiments.

### UR241-2 inhibits IL-1β-stimulated phosphorylation of IRAK4-long-form in multiple EOC cell lines

Next, we determined whether UR241-2 suppresses IL1β-induced IRAK4 phosphorylation. UR241-2 (20–150 nM) treatment for 4 hours prior to stimulation with mIL-1β (10 ng) successfully inhibited p-IRAK4 in HGS-3 and HGS-1 murine EOC cells (Fig-4A, middle and left). Next, we treated HCH-1 human EOC cells, with UR241-2 (5 and 15nM) for 4 hours prior to stimulation with hIL-1β (10 ng) for 45 minutes. UR241-2 treatment suppressed IL-1β induced IRAK4 phosphorylation (Fig-4A, right) Similarly,

**Figure-4:**
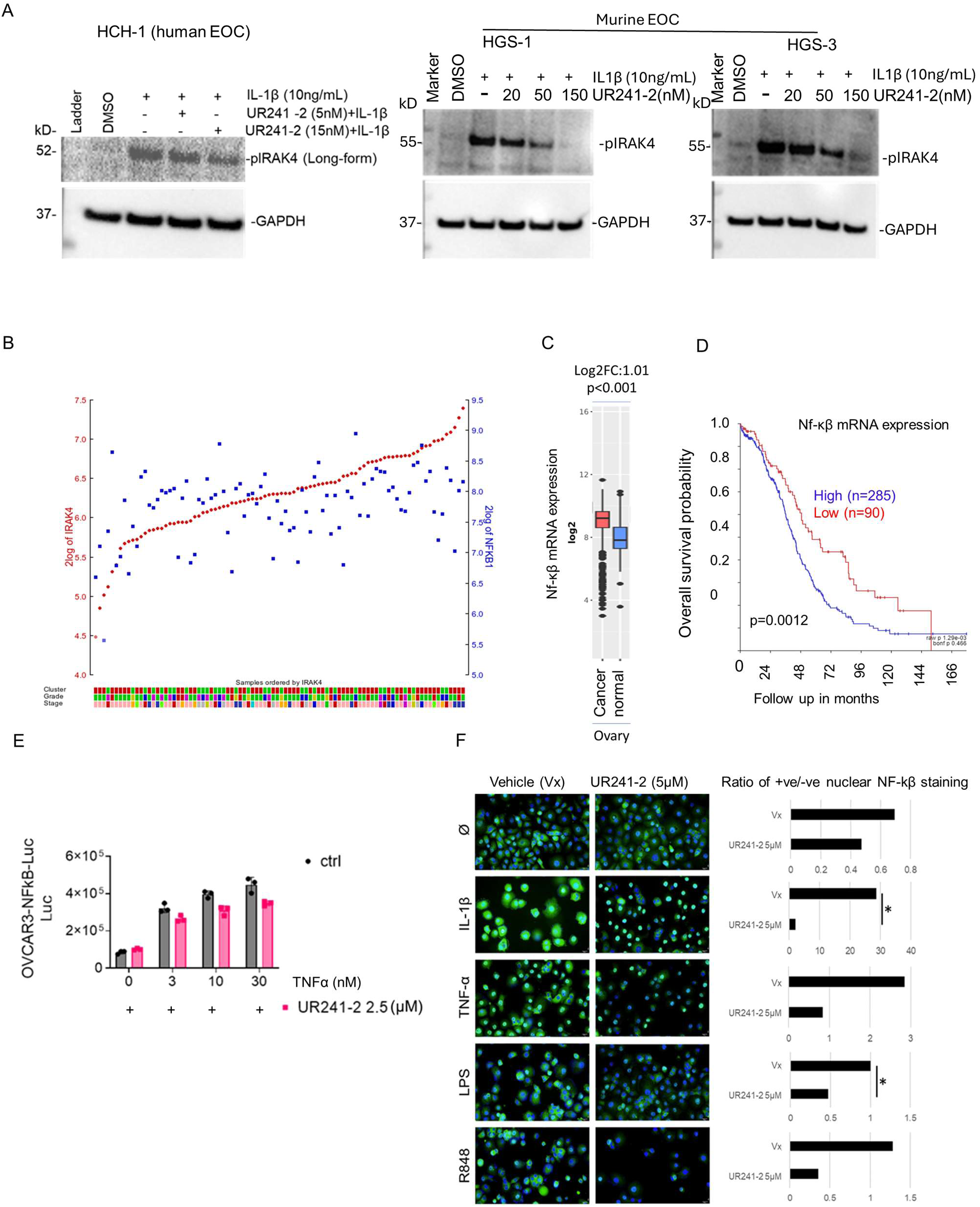
(**A**): UR241-2 (5-150nM) treatment for 4hrs blocks human and murine IL1β (5nM, 30 minutes) induced IRAK4-phosphorylation in both human (HCH-1) and murine high-grade serous EOC cells (HGS-1 and -3) cells. (**B**): Analysis of ovarian cancer patient’s microarray database (Kim_161-DESeq2_vst-ensh38e100) using R2 Genomics Analysis and Visualization Platform tools showed strong correlation of IRAK4 mRNA expression with NF-κβ mRNA expression in ovarian tumors (R=0.581, p=6.29e^-016^). Another database (TCGA-381-tpm-gencode36) showed similar correlation (R=0.523, p=5.28e^-018^) between IRAK4 and NF-κβ mRNA expression levels in ovarian tumors. (**C**): Analysis of ovarian cancer patient’s microarray showed using the GENT2 software showed that NF-κβ mRNA is overexpressed (Log2-Fold-change Log2FC=1.1) in malignant ovaries than normal (p<0.001). (**D**): NF-κβ mRNA overexpression showed significant risk of mortality among EOC patients (p=0.0012). The EOC patient’s microarray data was analyzed using the R2-Genomics Analysis and Visualization Platform tools. (**E-upper**): UR241-2 (2.5μM) treatment reduced NF-κβ-luciferase reporter activity in stably transfected OVCAR-3-NF-κβ-Luc cell-lines. **(E-lower**): UR241-2 (2.5μM) treatment reduced NF-κB luciferase reporter activity induced by human TNF-α (3, 10 and 30nM) in stably transfected OVCAR-3-NF-κβ-Luc cell-lines. (**F**): UR241-2 (2.5μM) treatment, IL-1β (5ng), TNF-α (10ng), LPS (100nM) and R848 (TLR-7/8 agonist, 10μM). Ratio of positive/negative NF-κB nuclei are shown. NF-κβ (Green) nuclear migration was inhibited significantly in IL-1β and LPS stimulated cells. The number of NF-κβ nuclear positive cells were counted using ImageJ software. * indicates <0.05 (Student T-test).

### UR241-2 inhibits IL-1β-, TNF-α-, LPS-, and TLR-agonist R848-induced NF-κβ activity in EOC cells

To evaluate the effects of UR241-2 on NF-κB signaling, a key downstream node of the IL-1/TLR/MyD88/IRAK1/4 axis, we investigated its response to IL-1β, TNF-α, LPS, and the TLR7/8 agonist R848. NF-κβ overexpression has been linked to chemoresistance, cancer stem cell maintenance, metastasis, and immune evasion^5–7^. Inhibiting NF-κβ not only validates the pharmacologic mechanism of UR241-2 but also provides a strategy to counteract tumorigenic signaling by these stimuli. Analysis of EOC patient tumor microarray data revealed a strong correlation between IRAK4 and NF-κB mRNA expression (R=0.581, p=6.29e-16, database ID: Kim-161-Deseq2_vst-ensh38100) (Fig-4B). Further, analysis using the ZENT2 database showed that NF-κB mRNA expression was significantly upregulated (Log2Fold Change: 1.01, p < 0.001) in malignant EOC cells compared to normal ovarian cells (Fig-4C). Additionally, NF-κB overexpression was associated with poor prognosis in EOC patients (p = 0.0012) (Fig-4D). Given the role of NF-κB in ovarian tumorigenesis, we assessed the effect of UR241-2 on NF-κβ reporter activity in OVCAR-3 cells stably expressing NF-κβ-Luc (provided by Annunziata Laboratory^46^). Treatment with UR241-2 (2.5 µM) reduced NF-κβ reporter activity compared to DMSO (Fig-4E, upper). Additionally, UR241-2 (2.5 µM) reduced NF-κβ reporter activity following TNF-α stimulation (3, 10, and 30 nM) (Fig-4E, lower). Next, we investigated whether UR241-2 (5 µM) could block nuclear translocation of NF-κβ induced by IL-1β, TNF-α, LPS, and R848. As shown in Fig-4F, UR241-2 treatment significantly inhibited nuclear NF-κβ translocation in OVCAR-3 cells. Specifically, UR241-2 (5 µM) significantly reduced the nuclear NF-κβ (positive/negative expression ratio) compared to the vehicle control upon IL-1β and LPS stimulation (Fig-4F, left). Similarly, UR241-2 (5µM) reduced the +ve/-ve nuclear NF-κβ expression ratio in response to TNF-α and R848 (Fig-4F, right).

### UR241-2 inhibits EOC colony formation, proliferation, and cell division while suppressing SKOV-3 xenograft growth without affecting body weight or hematology in mice

To assess the impact of UR241-2 on EOC colony formation, we treated SKOV-3, OVCAR-3, and OVCAR-8 cells with increasing concentrations of UR241-2. Colony size was significantly reduced compared to the control in SKOV-3 at 5 µM (p = 0.0131), 10 µM (p = 0.0437), and 20 µM (p = 0.0077) (Fig-5A, upper). In OVCAR-3, 5 µM treatment significantly reduced colony size (p = 0.0488), while 10 µM (p < 0.0001 vs. control, p = 0.0142 vs. 5 µM) and 20 µM (p < 0.0001 vs. control, p = 0.0104 vs. 5 µM) exhibited greater suppression (Fig-5A, middle). Similarly, in OVCAR-8, 5 µM significantly reduced colony size (p = 0.0038), with 10 µM (p = 0.0012 vs. control, p = 0.0422 vs. 5 µM) and 20 µM (p = 0.0055 vs. control, p = 0.0116 vs. 5 µM) further reducing colony formation (Fig-5A, lower). The average colony size across treatment groups is summarized in Fig-5B. Next, we evaluated the effect of UR241-2 on EOC cell proliferation using the SRB assay^47^, which measures protein synthesis as an indicator of viability. Treatment with UR241-2 (20–60 µM) for 48 hours resulted in a dose-dependent reduction in viability across a panel of EOC cell lines, including ES2, HCH1, SKOV-3, OVCAR-3, and OVCAR-8 (Fig-5C). To determine whether UR241-2 affects cell division, we assessed Histone-H3 phosphorylation, a marker of mitotic activity. Treatment with UR241-2 significantly reduced the percentage of actively dividing cells in both OVCAR-3 (p = 0.0024 at 20 µM) and SKOV-3 (p = 0.0047 at 10 µM, p = 0.0003 at 20 µM) (Fig-5D).

**Figure-5:**
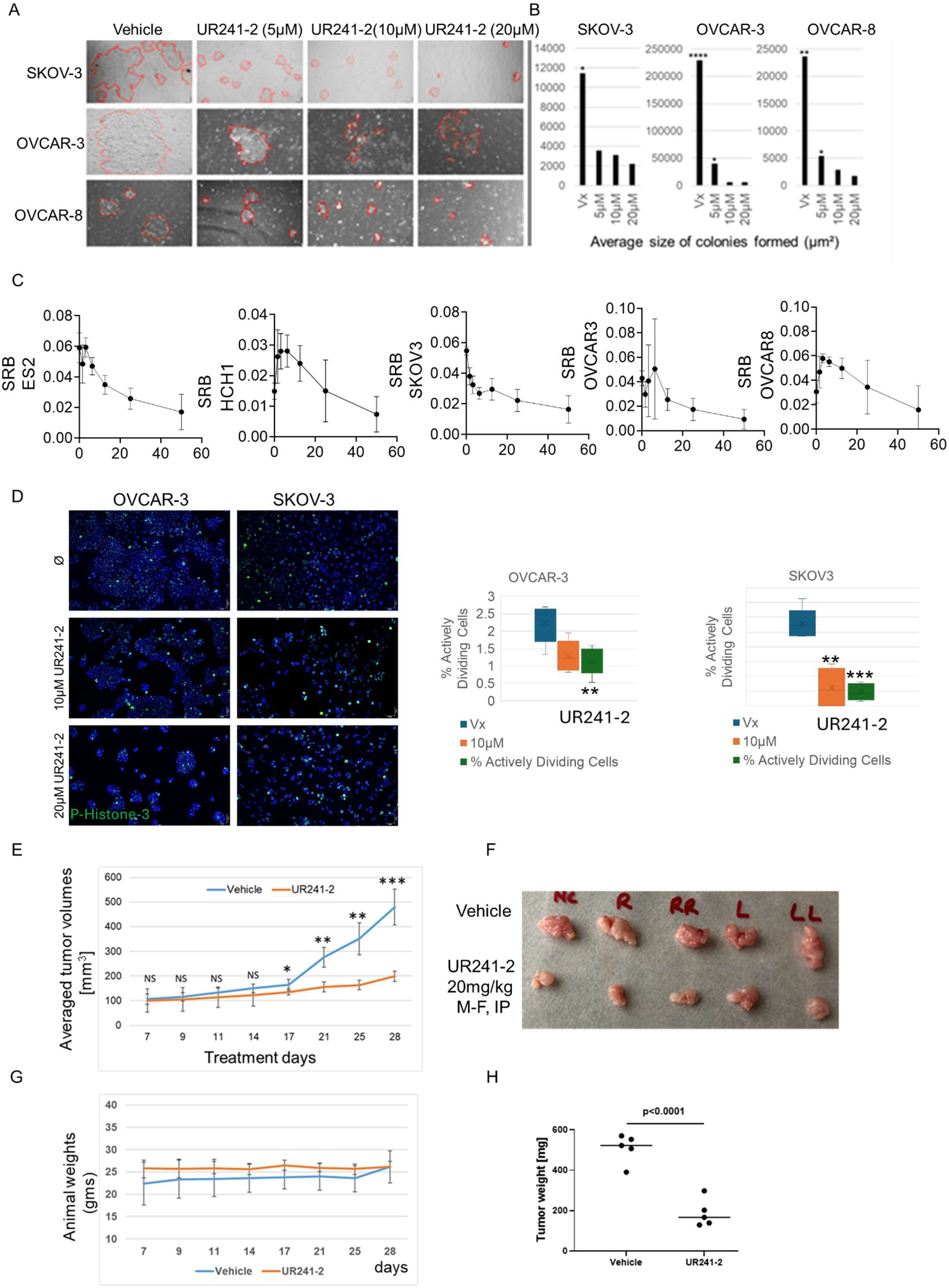
(**A**): UR241-2 (5, 10 and 20 µM) treatment for 7 days blocks colonies formed from 2000 cells seeded in a 6 well plate compared to control. One representative images of multiple images of each well that were collected are shown. (**B**): Average size of colonies in vehicle versus 5, 10 and 20 μM groups were analyzed by ImageJ software. Non-parametric T-test was conducted to determine the statistical differences between the control and various treatment groups. * indicates <0.05 (Student T-test). (**C**): UR241-2 (20-60μM) treatment reduced cell viability in ES2, HCH1, SKOV-3, OVCAR-8 and OVCAR-3 EOC cells. The cell viability was measured by SRB assay which measures the total protein synthesis in drug treated cells compared to control vehicle. (**D**): UR241-2 (10-20μM) treatment reduced cell division in SKOV-3 and OVCAR-3 cells. The dividing cells were captured by pHistone-3 staining. SKOV-3 and OVCAR-3 cells (10,000/well) were seeded in an 8-well EasyCell chamber slides overnight. The cells were treated with vehicle and UR241-2 ((10-20μM). Cells were fixed after 24-hours using neutral buffered formalin solution (35µL) for 20 minutes. The media was removed, and cells were washed repeatedly with TBST (500µL/5 min). Fixed cells were stained with p-Histone-H3 antibody (Cell Signaling Technology, cat#9701, 1:500 dilution, overnight) and then counterstained with Dylight-488(Vector laboratories, cat#DI-1488) secondary antibody (1:2000). The cells were washed repeatedly with TBST (500µL/5 min). Washed cells were stained again with the Vectashield mounting media containing DAPI (Vector Lab, cat#H-1200-10), protected with a cover glass slide, then imaged on an Olympus BX41 microscope. All images taken with through a 10x ocular and 20x objective. All cells in each field were manually counted and assessed as either actively dividing (metaphase, anaphase, telophase) or not. OVCAR-3 vehicle group had significantly more actively dividing cells than the 20μM treatment group (p=0.0024). SKOV3 vehicle group had significantly more actively dividing cells than both 10μM and 20μM treatment groups (**p=0.0047, ***p=0.0003). (**E**): UR241-2 treatment (20mg, M-F, IP) reduced the growth of SKOV-3 cells derived xenografts growing in NSG mice. 10 NSG mice implanted subcutaneously with SKOV-3 cells (1 million/mice in DMEM+Matrigel (1:1, 100µL/mice) were randomized and treated for 28-days. Tumor sizes were measured on the days indicated in the X-axis. Mice were euthanized and tumors and peripheral blood were harvested. Averaged tumor volumes differed significantly between the control and treatment groups. *=p<0.05, **=p<0.005, ***=p<0.0005. (**F**): Images of the tumors formed in vehicle and UR241-2 treated mice are shown. (**G**): Animal weights did not differ. (**H**): Tumor weights between vehicle and treated groups differed significantly. The tumor weights of the individual mice in the control and treatment groups were plotted using GraphPrism Version 8.0. *p*=0.0001.

Given these findings, we next examined the *in vivo* efficacy of UR241-2 using a SKOV-3 xenograft model. Based on pharmacokinetic data (Supplementary Fig-3G), we administered UR241-2 at 20 mg/kg (IP, M–F, once daily). As shown in Fig-5E, UR241-2 treatment significantly inhibited tumor growth starting on Day 17 (p < 0.05). By Day 28, tumors in the UR241-2-treated group were significantly smaller than those in the vehicle group (p < 0.0005). Notably, UR241-2-treated mice did not exhibit weight loss or signs of poor well-being (Fig-5G). After euthanasia, tumors were harvested, imaged, and weighed. Tumor weights in UR241-2-treated mice were significantly lower than in vehicle-treated mice (p < 0.0001) (Fig-5F, H). An estimation plot further confirmed that tumor sizes were significantly different between treatment and control groups (Supplementary Fig-4A). Blood chemistry of peripheral blood collected prior to euthanasia showed that hemoglobin (HB) levels did not vary between the control and UR241-2 treated mice (Supplementary Fig-4B).

### UR241-2 inhibits tumor development at injury sites in syngeneic high-grade serous murine EOC models and modulates inflammatory immune cell populations

To evaluate the anti-tumor effects of UR241-2, we utilized a syngeneic MIM EOC model that recapitulates cell adhesion at the site of injury. HGS-3 cells^48^, when implanted intraperitoneally (IP), also mimic the fallopian tube origin of EOC, a new line of thinking in the origin of EOC^49^. UR241-2 treatment significantly reduced tumor size at the injury site (Fig-6A) but did not affect omental tumor burden (Fig-6B), suggesting that injury- or inflammation-driven signaling is necessary for IRAK4-targeted interventions. To assess immune modulation by UR241-2, leukocytes were isolated from either peritoneal lavage or tumor tissue of HGS-3 tumor-bearing mice and analyzed by flow cytometry (Fig-6C). Representative flow cytometry plots of F4/80 and MHCII-expressing myeloid cells from the peritoneal lavage of vehicle (Vx)- and UR241-2-treated mice are shown. UR241-2 treatment significantly increased both the percentage and number of MHCII^hi^ macrophages in the peritoneal lavage compared to the vehicle group (Fig-6D, left). Additionally, UR241-2 treatment reduced the proportion of MHCII^Low^ macrophages (Fig-6D, right), indicating a shift from a more anti-inflammatory environment toward a pro-inflammatory state enriched with MHCII^hi^ antigen-presenting cells. We also observed a significant decrease in the percentage of F4/80^hi^CD206^+^ M2-like macrophages within the peritoneal cavity of tumor-bearing mice treated with UR241-2 (Fig. 6E). Furthermore, UR241-2 treatment significantly increased neutrophil populations in both tumors and the peritoneal cavity (Fig-6F, G). Given that neutrophils exhibit functional plasticity with both pro- and anti-tumor roles, we quantified MHCII-expressing neutrophils. UR241-2-treated mice exhibited a higher number of MHCII^+^ neutrophils in both peritoneal lavage and tumors, suggesting a pro-inflammatory, anti-tumor phenotype (Fig-6F, G). To determine whether IL-1R1 loss mimics UR241-2-mediated immune modulation, we examined *Il1r1*^KO^ HGS-3 tumor-bearing mice. Unlike UR241-2 treatment, *Il1r1*^KO^ mice did not exhibit similar alterations in myeloid populations (Supplementary Fig-5). This suggests that IL-1R1 signaling is required to initiate inflammatory responses in the peritoneal cavity and tumor microenvironment, while IRAK4 inhibition by UR241-2 drives a more pronounced pro-inflammatory immune shift in this context, marked by the increase in MHCII expressing cells and decrease in macrophages with the typical M2 marker CD206.

**Figure-6:**
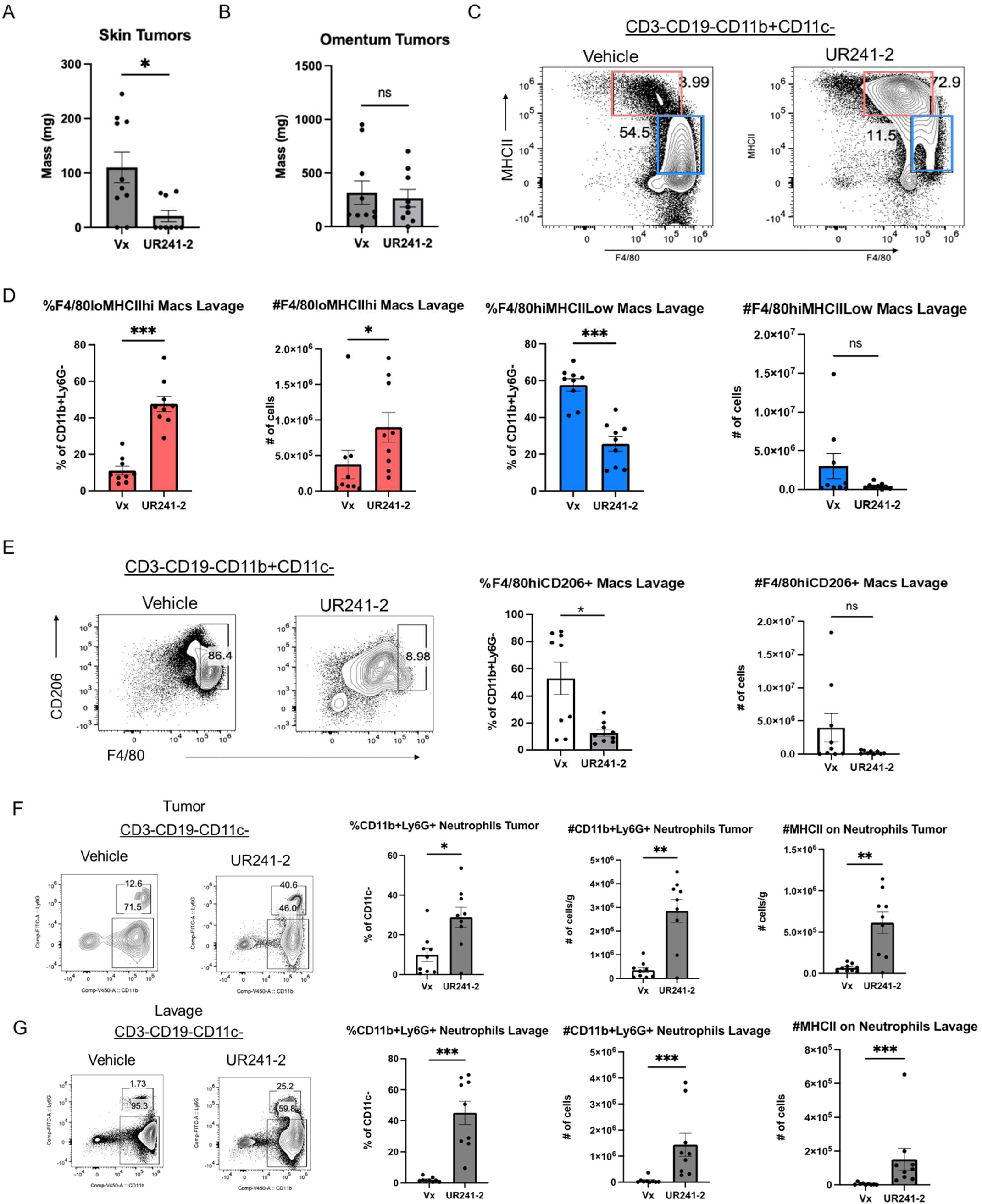
IRAK4 inhibition via UR241-2 decreases tumor burden at the needle injury site and increases the number of MHCII expressing myeloid cells in the peritoneal cavity and tumor. (**A**): tumor weights on the needle injury site were significantly lower in the UR241-2 treatment group than vehicle treated animals. Statistical significance was determined using Mann-Whitney test, *p < 0.033, **p < 0.002, ***p < 0.001. (**B**): Tumor weights on omentum did not differ between UR241-2 treatment group than vehicle treated animals (NS). Two of the three replicates are combined and shown. ***IRAK4 inhibition via UR241-2 increases the number of MHCII expressing myeloid cells and decreases MHCII low and CD206+ macrophages in the peritoneal cavity***. Leukocytes were isolated from the peritoneal lavage or tumor of HGS-3-tumor bearing mice and stained for flow cytometry. (**C**) Representative flow cytometry plots of F4/80^lo^MHCII^hi^ M1 macrophages and F4/80^hi^MHCII^Low^ macrophages from the peritoneal lavage of HGS-3 tumor bearing mice treated with Vehicle or UR241-2. (**D**) Percentage and cell number of F4/80^lo^MHCII^hi^ M1 macrophages and F4/80^hi^MHCII^Low^ macrophages from C. **(E)** Representative flow cytometry plots of F4/80^hi^CD206^+^ macrophages from the peritoneal lavage of HGS-3 tumor bearing mice treated with Vehicle (Vx) or UR241-2. (**E-right**) Percentage and cell number of F4/80^hi^CD206^+^ macrophages. ***IRAK4 inhibition reduces neutrophil numbers in the peritoneal cavity and tumors of HGS3 bearing mic*e: (F**): Flow cytometry plots showing CD11b^+^Ly6G^+^ neutrophils and MHCII^+^ neutrophils in the HGS-3 tumor. Percentage and cell number per gram are quantified to the right. (**G**): Flow cytometry plots showing CD11b^+^Ly6G^+^ and MHCII^+^ neutrophils in the peritoneal lavage of HGS-3 tumor bearing mice. Percentage and cell number are quantified to the right. N= 9 mice per group from two independent experiments.

### Bulk RNA-Sequencing analysis of UR241-2-treated tumors reveals changes in ECM organization and neutrophil-related gene expression that corroborate with flow cytometry

Bulk RNA-sequencing (bulk-seq) analysis of UR241-2-treated tumors revealed significant alterations in gene expression associated with extracellular matrix (ECM) organization and neutrophil-mediated immunity. Genes from tumors isolated in Fig-6A were processed and analyzed. Principal component analysis (PCA) (Fig-7A) and volcano plots (Fig-7B, 7C) illustrate differences between vehicle- and UR241-2-treated tumors. Among the top 30 significantly downregulated pathways (adjusted p-values), ECM organization genes, which play a crucial role in tumor cell adhesion, were notably suppressed (Fig-7D). Conversely, neutrophil-associated pathways, including neutrophil-mediated immunity, neutrophil activation in immune response, were significantly upregulated in UR241-2-treated tumors (Fig-7E). Heatmaps of UR241-2-treated versus control tumors depict global gene expression changes (Fig-7F) and ECM gene downregulation (Fig-7G). Functionally, UR241-2 treatment reduced HGS-3 cell adhesion to a collagen basement membrane in a dose-dependent manner (Fig-7H). Additionally, HGS-3 cells exhibited significantly reduced migration *in vitro* at UR241-2 concentrations as low as 250 nM (Fig-7I). Taken together, this data indicates that UR241-2 treatment leads to downregulation in ECM organization genes, and upregulation in neutrophil-activation gene pathways, correlating with *in vitro* cell adhesion and wound healing data and *in vivo* flow cytometry data respectively. This provides some mechanistic indication to how IRAK4 inhibition could lead to a decrease in tumor cell invasion at the site of injury, as well as increased anti-tumor inflammatory responses in the peritoneal cavity of tumor bearing mice.

**Figure-7:**
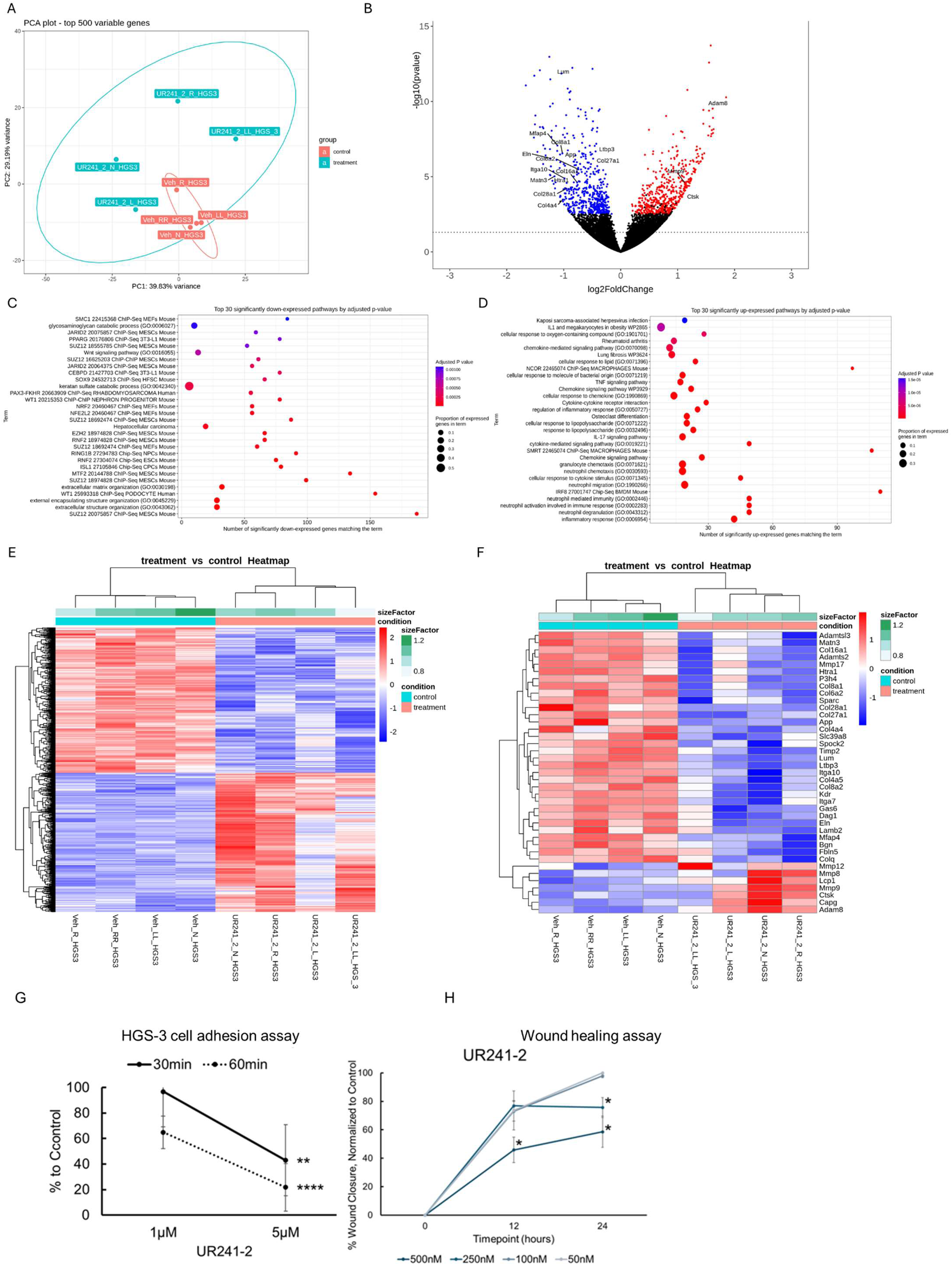
(**A**): PCA plot of Bulk-Seq analysis of control and UR241-2 treated tumors. (**B**): Volcano plot of genes overexpressed and downregulated between control and treatment groups are shown. Downregulated genes belonging to ECM family are shown. (**C**): Gene-families downregulated by UR241-2 are shown. (**D**): Gene-families upregulated by UR241-2 are shown. (**E**): Heatmap of the genes altered between control and treatment. (**F**): Heatmap of the ECM family of genes downregulated by UR241-2 are shown. (**G**): HGS-3 cells treated with UR241-2 demonstrated an inhibited ability to adhere to collagen compared to control, with cells treated at 5μM showing significantly less adhesion than the 1μM group at both the sixty (p<0.00005) and thirty (p=0.00068) minute timepoints. (**H**): Cell migration in HGS-3 cells treated with 500nM UR241-2 were significantly inhibited compared to control in just 12 hours (p<0.02), and both the 250nM and 500nM groups showed a significantly greater impact than the 100nM, 50nM, and control treatment groups at 24 hours (p<0.05).

## Materials and Methods

### Microarray database analysis

Kaplan-Meier survival outcomes among WOC patients (**Fig-1A, 1B, 2B, 2C**, **2D and 4D**) were analyzed using the databases (tcga-381-tpm-gencode36) and tools available at R2-Genomics and Visualization Platform (https://hgserver1.amc.nl/) using the system selected gene expression cut-off points. For determining the expression of IRAK4 the database (Mori-171-rma_sketch_hugene10t) was also analyzed. Expression of IRAK4 in venous-invasion or no-invasion was examined in the database (TCG-541-custom-tcgaovag1) (**Fig-2G**). Gent2 database (http://gent2.appex.kr/gent2/) was used to compare the IRAK4 (**Fig-2E**) and NF-κβ expression (**Fig-4C**) differences between normal ovaries versus malignant ovary. Gent2 database was also used to compare the IRAK4 expression differences between various EOC stages (date accessed: 10/2/2023) (**Fig-2F**). To determine the correlation between IRAK4 and NF-κβ gene expressions, the EOC microarray database (Kim-161-DESeq2_vst-ensh38e100) were also analyzed using R2-Genomics and Visualization Platform using the inbuilt tools (**Fig-4B**).

### MiM model

3-7 million HGS-3 murine high-grade serous EOC cells suspended in serum and antibiotic free DMEM medium were injected intraperitoneally in C57/BL6 mice or their derivative IL1R1^KO^, IL1RN^KO^ and NLRP3^KO^ mice using 21G needles. Mice are monitored for 40-60 days when most of the mice will present tumor nodules at the site of needle injury. Skin of the mice are shaved carefully to remove the fur till the tumors become palpable.

### EOC tumor burden in syngeneic Il1r1^KO^, Il1rn^KO^ and Nlrp3^KO^ mice

*Il1r1*^KO^ (B6.129S7-Il1r1tm1Imx/J), *Il1rn*^KO^ (**Fig-1E**) (B6.129S-Il1rn tm1Dih/J) (**Fig-1K**) and *Nlrp3*^KO^ (Strain #:021302 - B6.129S6-Nlrp3tm1Bhk/J) mice were bred in-house from the breeders obtained from Jax laboratories. Homozygous *Il1r1*^KO^ (n=5), *Il1rn*^KO^ (N=5) and *Nlrp3*^KO^ (n=5) mice were implanted with HGS-3 mouse HGS-3 EOC cells (3-4.5 million/mice) intraperitoneally. Similarly, WT C57BL/6 mice (n=5) were also injected with HGS-3 murine HGS EOC cells (3-4.5 million/mice). After 45 days, mice were bled via mandibular vein and ∼25µL of blood collected for complete blood counts (CBC) was analyzed in EDTA-tubes using ElementHTS+ system. Mice were euthanized soon after and photographed to record the tumors formed at the injury sites. Spleen, omentum, and peritoneal lavage were collected from each mouse. Omentum weights were recorded and weighed. Spleen was crushed in serum-free RPMI1640 medium (Gibco, cat#: 22400-089) and filtered. Tumors/omentum were processed similarly. Harvested cells were broken down into single cell suspension and analyzed by flow cytometry following the conditions described below.

### Flow cytometry

Omentum and tumors were minced in plain RPMI and incubated in collagenase (2mg/ml), DNase (25ug/ml), and hyaluronidase (0.1mg/ml) for 30min-1hr at 37^0^C. The suspension was strained through 70µm strainers, diluted with 10ml of cold complete RPMI (10% FBS) and centrifuged at 300 x g for 5 mins at 4^0^C. The cells were resuspended in PBS for cell counting and stained for flow cytometry. The spleen was isolated and crushed through a 70µm strainer in cold complete RPMI (10% FBS) and transferred to tubes and the cells were centrifuged at 300 x g for 5 mins at 4^0^C. The resulting pellet was resuspended in 1X Red blood cell lysis buffer for 4-5mins at room temperature with occasional shaking. The suspension was diluted with complete RPMI (10% FBS) and centrifuged at 300 x g for 5 mins at 4^0^C. The pellet was resuspended in RPMI (10% FBS) for cell counting and stained for flow cytometry. The peritoneal lavage was collected using a 5ml syringe and 20G needle to inject 5ml of PBS into the peritoneal cavity. The abdomen was massaged for 10-15 seconds, and the peritoneal lavage was collected using the needle. The needle was removed, and the resulting suspension dispersed into a tube on ice. The suspension was centrifuged at 300 x g for 5 mins at 4°C and resuspended in 10ml of PBS and passed through a 70µm strainer. The suspension was centrifuged at 300 x g for 5 mins at 4^0^C and resuspended in PBS for cell counting and stained for flow cytometry. After staining, all tissues were fixed and analyzed using a Cytek Aurora Full Spectrum Flow Cytometer. The staining antibodies were used at a dilution of 1:200. The list of antibodies, catalog numbers and source are described in the Supplementary list-6.

### Characterization data of UR241-2 and analogs

The synthesis of UR241-2 and analogs (PSP-099 and PSP-100) (**Fig-2I**) was achieved in milli gram scale. The compounds were characterized by ^1^H- and ^13^C-NMR and Mass spectrometry (MS) Supplementary data-7. Purity was established by HPLC. The compounds tested in this study were each more than 97% purity by HPLC.

### Hotspot Kinase, Global kinome analysis and cell based Nanobret kinase binding assay

UR241-2 and analogs along with controls PF-0665083 and CA4948 (**Fig-2I**) were screened for the IRAK1, -2, and -4 kinase activity in 10-dose singlet curve mode using the Hotspot kinase assay offered by Reaction Biology Inc. (**Fig-2K**). Selectivity of UR241-2 was analyzed against more than 680 kinases in a Global kinome screening assay offered by Reaction Biology Inc at 3 escalated doses. Dendrograms of kinase activities at 50 and 500nM dose of UR241-2 were prepared using the KinMap^50^ (http://www.kinhub.org/kinmap/) (**Fig-2L**). The specificity of UR241-2 against IRAK1, IRAK4, LRRK2, FLT3, MAP4K1, MAP4K2, MAP4K3, MAP4K5. The kinases that showed comparable kinase inhibition in Hotspot kinase screening *in vitro* were determined using HEK293-cell-based assays (**Fig-2M**).

### In silico docking and molecular docking simulations

The crystal structure of IRAK4 was downloaded as PDB from the PDB bank (RCSB ID: 2NRU). The protein was prepared prior to docking, which involved selecting a single chain out of the four identical ones that were present. Further preparation was handled by the GNINA docking suite^51^ (https://github.com/gnina) with machine learning (ML) hosted on Google Colab. Next, the docking grid box was created to serve as the search area for the GNINA docking suite. The T12 ligand was selected as the center of the grid box. This area is the ATP binding site and was chosen due to the design of the molecule serving as a competitive inhibitor, completing protein preparation. Ligand preparation began conversion from SMILES to SDF. The SDF file was uploaded to Google drive. Further preparation of the ligand continued with the protonation of the ligand and the subsequent geometry optimization using Torch ANI, with the convergence threshold set to 0.0001. This completed the ligand preparation. Next, we selected our docking parameters for GNINA. Exhaustiveness was set to 64, buffer space was set to 4 angstroms, and refinement scoring was used. Then we performed docking via the GNINA docking suite and obtained multiple poses. The best pose was selected based on the highest affinity and the predicted location of our ligand (**Fig: 3A-C**). Ligands outside of the ATP binding domain were disregarded. This completed the docking phase. The docked ligand and protein were then moved to undergo simulations. Simulations were done using the OpenMM engine. A notebook made by the “Making it rain” team had graciously written the code for the engine on Google Colab. Parameters for topology generation were set as following: the NaCl concentration was set 0.15 mol, padding distance was set to 4, the force field used was the AMBER99SB, the water type model was set to SPC/E, pH was set to 7.4, and finally gaff-2.11 was used to generate ligand force field. MD equilibration was done prior to the MD simulation. Both were simulated at a temperature of 298K and 1-bar pressure. The equilibration process was conducted for 0.5 nanoseconds and the simulation was conducted for 5 nanoseconds. Images were generated by the libraries within the notebook (**Fig:3D-I**). Physicochemical properties of the compounds were calculated with ChemDraw software using the chemical analysis tools (**Fig-2J**).

### Effects of UR241-2 on NF-κB activity in EOC cells

OVCAR-3 cells (20,000/well) were seeded in 8-well chambers slides and cultured in DMEM (Gibco, cat#: 11885-084) supplemented with 10% FBS (Avantor, cat# 89510-186), 1% penicillin/streptomycin (Gibco, cat#:15140-122). Media was replaced with serum free DMEM media cells were stimulated with either IL1β (5ng), TNF-α (5ng), LPS (5µg), R848 (10µM), or negative control, for 24 hours prior to treatment start. Each group was either treated with 5μM UR241-2 or DMSO vehicle for 24 hours prior to analysis. Cells were then fixed with neutral formalin for 25 minutes at 4°C and washed with PBST(3×500µL). The cells were stained with NF-κB p65 (D14E12) XP® Rabbit mAb primary antibody (Cell Signaling Tech, cat#:8242) in PBST (1:250) overnight. TBST containing antibody media was removed, washed with 2×500uL TBST/5 minutes each, and stained with DyLight488 anti-rabbit IgG (DI-1488, Vector Laboratories) (1:2000, 1 hours under aluminum foil wrap). After an hour, antibody was removed, washed with 5×500uL TBST, 15 minutes each and finally EZSlide was dismantled. Resulting glass slides was counterstained with Vectashield Antifade Mounting Medium with DAPI (Vector Laboratories, cat#: EW-93952-24), covered with glass coverslips (VWR, cat#48393-081), and stored in a slide-holder packet at 4°C/dark until examined under microscope. Epifluorescent images taken on an Olympus BX41 microscope with Olympus DP74 camera using CellCens software under 20x objective and 10x ocular magnification (200x). Cells manually counted and assessed as follows: Cells with clear nuclear NF-κβ staining greater than control were counted as “positive”, any other cell was counted as “negative” regardless of signal intensity outside of nucleus.

### Western blot analysis

Human (HCH-1) or murine EOC (HGS-1 and HGS-3) cell-lines were seeded in 60mm^3^ (Nest, cat#7050010 overnight in complete DMEM media. Media in 60mm^3^ dishes were replaced with serum free DMEM and stimulated with human IL1β (10ng, 30 minutes) or murine IL1β (10ng, 30 minutes) without or with pretreatment with UR241-2 (HCH-1, 5-15nM, 4 hrs). For murine EOC cell-lines HGS-1/3, 20, 50 and 150nMoles of UR241-2 were used (**Fig-4A**). The dishes containing the HCH-1 and HGS-1/3 cells were washed with PBS and lysed with 120uL 1X cell-lysis buffer (Cell Lysis Buffer (10X), Cell Signaling Tech, cat #9803). The lysate was collected in 1.5 ml tube and incubated on ice for 10 min. The lysates were then centrifuged at 14,000 rpm for 10 min at 4°C for removing cell debris. The lysates were electrophoresed using NuPage 4-12% Bis-Tris gels (Invitrogen, cat#NP0322BOX) and transferred semi-dry to PVDF membrane cat#1703966) were all incubated in NuPAGE transfer buffer for 10 min before assembly. The transfer was done at room temperature for 90 minutes at 120 V constant voltage using the PVDF membrane (BioRad, cat#1620177) and filter pads (Bio-Rad, cat#1703969). The PVDF membranes were washed with methanol and then blocked with 5% fat free milk. Chemiluminescent detection was done with a Super Signal West FemtoLuninol/Enhancer (ThermoScientific, cat#1859022 and peroxide cat#1859023) or using Amersham ECL Prime (peroxide solution, cat#RPN2232V2 and enhancer cat#29018903; Cytiva) and probed with p-IRAK4 antibodies (Cell Signaling, cat#11927) (dilution: 1:500). Membranes were imaged using BioRad Chemidoc MP imaging system.

### Colony Formation Assay (Fig-5A)

OVCAR-3, OVCAR-8, and SKOV-3 cell lines (20,000/well) were seeded in sterile 8-well glass Millicell EZSlide chambers (Merck Millipore; cat#: PEZGS0816) and cultured in DMEM (Gibco, cat#: 11885-084) supplemented with 10% FBS (Avantor, cat# 89510-186), 1% penicillin/streptomycin (Gibco, cat#:15140-122). Cells were treated with either 5μM, 10μM, or 20μM of UR241-2, or DMSO vehicle for 7 days prior to assessment of colony formation. OVCAR-3 colonies were monitored for 14 days. Phase-contrast images were taken on an Olympus BX41 microscope with Olympus DP74 camera using CellCens software under 20x objective and 10x ocular magnification (200x). Colony borders manually traced in Fiji ImageJ software and the area was measured in μm². Differences between control and treatment groups were analyzed using T-test.

### Cell proliferation assay (Sulforhodamine B Assay, Fig-5C)

OVCAR-3, OVCAR-8, SKOV-3, ES2, and HCH1 human EOC cells (5000/well) were cultured in flat-bottom 96-well plates with 100μL of DMEM (Gibco, cat#: 11885-084) supplemented with 10% FBS (Avantor, cat# 89510-186), 1% penicillin/streptomycin (Gibco, cat#: 15140-122) until ∼50% cell density was met. Cells were then treated for 48 hours with a series of 2-fold dilutions of UR241-2 resulting in the concentrations of 50μM, 25μM, 12.5μM, 6.25μM, 3.12μM, and 1.56μM, as well as DMSO vehicle and negative controls. Cells were fixed with the addition of 70μL 20% TCA and incubated for 1 hour at 4°C. The TCA solution was aspirated, and the plate gently washed once with 200μL dH₂O per well. 100μL of 1x SRB solution diluted in non-supplemented RPMI1640 was added to each well and incubated for 15 minutes at room temperature and protected from light. Solution was aspirated, and plate washed with 200μL 1% acetic acid for 5 minutes on a plate rocker before aspirating off solution. This wash was repeated twice again, and the plate allowed to sit uncovered for residual acetic acid solution to evaporate. 100μL of 10mM Tris base solution was added to each well and the plate was incubated at room temperature for 10 minutes, rocking gently and protected from light. Plate was tapped gently to homogenize dye solution in wells, and then read for absorbance at 510nm using the BioTek Synergy 2 plate reader.

### Cell division analyses (Fig-5D)

OVCAR-3 and SKOV-3 cells (10,000 cells/well) were seeded in sterile 8-well glass Millicell EZSlide chambers (Merck Millipore; cat#: PEZGS0816) and cultured in DMEM (Gibco, cat#: 11885-084) supplemented with 10% FBS (Avantor# 89510-186), 1% penicillin/streptomycin (Gibco#: 15140-122). Cells were treated with either 10μM or 20μM of UR241-2, or DMSO vehicle for 48 hours prior to assessment. Cells were then fixed with neutral formalin solution for 25 minutes at 4°C, washed with TBST (3×500mL, 5 minutes each) and stained with p-Histone H2A primary antibody (Cell Signaling Tech, cat#:9718p, 1:1000 in TBST) and incubated overnight. TBST containing antibody was removed, washed 2×500uL TBST 2 times/5 minutes each and stained with DyLight488 anti-rabbit IgG (DI-1488, Vector Laboratories) (1:2000, 1 hours under aluminum foil wrap). After an hour, antibody was removed, washed with 5×500uL TBST, 15 minutes each, and finally EZSlide was dismantled. Resulting glass slides was counterstained with Vectashield Antifade Mounting Medium with DAPI (Vector Laboratories, cat#: EW-93952-24) and covered with glass cover-slides (VWR, cat#48393-081) and stored in a slide-holder packet at 4°C/dark till examined under microscope. Epifluorescent images taken using an Olympus BX41 microscope with Olympus DP74 camera using CellCens software under 20x objective and 10x ocular magnification (200x). Cells manually counted and assessed as follows: cells visibly replicating (metaphase, anaphase) were manually counted and assessed as a percent of the total cells in the field. Cells not totally within the border of the image were not counted.

### Xenograft model (Fig-5E)

SKOV-3 cells (1million cells/mice) were implanted subcutaneously in NSG mice in cold serum free DMEM:Matrigel (1:1). After a week, mice with palpable tumors were randomized and treated with either vehicle (200uL/mice, IP, M-F) or UR241-2 (20mg/kg, IP, M-F). Tumor diameter (L) and width(w) was measured using a digital caliper. On 28^th^ day, mice were euthanized, tumors and peripheral blood from each mouse were collected. The tumors weight was measured using a calibrated balance and hematology of the blood was analyzed using ElementHT-5+ CBC machine.

### Hematology

25-30 µL peripheral blood was collected via retromandibular vein from mice in EDTA-tubes. Complete blood counts (CBC) were analyzed using the HT-5 CBC blood analyzer machine using the system set parameters.

### Effect of UR241-2 on murine EOC tumor burden in syngeneic C57BL/6 mice (Fig-6A-B)

C57BL/6 mice were bred in house from the breeders obtained from Jax laboratories or were purchased from the Jax. C57BL/6 mice (n=5 each for control and UR241-2) were implanted with HGS-3 mouse HGS-3 EOC cells (3-4.5 million/mice) intraperitoneally. Two days post tumor cell inoculation, mice were treated with either vehicle or UR241-2 (30mg/kg, 100µL, M-F, IP). DMSO solution of UR241-2 (1µL=200uL, 90mg in 450µL DMSO) was dissolved in formulation made of [H2O (300µL)+PEG6000 (40% in water, 150µL)+EtOH(150µL)] which generates a clear solution after short vortexing. Equivalent volume of the solvent was injected in each animal of the vehicle group. Treatment continued till 44^th^ day. After 45 days, mice were bled via mandibular vein and ∼25µL blood was collected for complete blood counts (CBC) and analyzed in EDTA-tubes using ElementHTS+ system. Mice were euthanized soon after and photographed to record the tumors formed at the injury sites. Spleen, omentum and lavages were collected from each mouse. Omentum and spleen weights were recorded and weighed. Spleen was crushed in serum free RPMI1640 medium (Gibco, cat#: 22400-089) and filtered. Tumors/omentum were also crushed similarly. Crushed cells were broken into single cell suspension and analyzed by flow cytometry following the conditions described below.

### Bulk-seq analysis of the tumors (Fig-7)

Total RNA was isolated using the RNeasy Plus Mini Kit (Qiagen, Valencia, CA) per manufacturers recommendations. The total RNA concentration was determined with the NanopDrop 1000 spectrophotometer (NanoDrop, Wilmington, DE) and RNA quality assessed with the Agilent Bioanalyzer (Agilent, Santa Clara, CA). The Illumina Stranded Total Sample Preparation Kit (Illumina, San Diego, CA) was used for next generation sequencing library construction per manufacturer’s protocols. Briefly, ribosomal-depletion was performed on 100ng total RNA followed by RNA fragmentation. First-strand cDNA synthesis was performed with random hexamer priming followed by second-strand cDNA synthesis using dUTP incorporation for strand marking. End repair and 3’adenylation was then performed on the double stranded cDNA. Illumina adaptors were ligated to both ends of the cDNA and amplified with PCR primers specific to the adaptor sequences to generate libraries of approximately 200-500bp in size. The amplified libraries were hybridized to the Illumina flow cell and sequenced using the NovaSeq6000 sequencer (Illumina, San Diego, CA). Paired-end reads of 150nt were generated and a read depth of ∼50M reads was targeted for each sample. Raw reads generated from the Illumina basecalls were demultiplexed using bcl2fastq version 2.19.0. Quality filtering and adapter removal are performed using FastP version 0.20.1 with the following parameters: “--length_required 35 --cut_front_window_size 1 -- cut_front_mean_quality 13 --cut_front --cut_tail_window_size 1 --cut_tail_mean_quality 13 --cut_tail -y -r”. Processed/cleaned reads were then mapped to the mouse reference genome GRCm39 using STAR_2.7.6a with the following parameters: “—twopass Mode Basic --runMode alignReads --outSAMtype BAM SortedByCoordinate – outSAMstrandField intronMotif --outFilterIntronMotifs RemoveNoncanonical –outReads UnmappedFastx”. Genelevel read quantification was derived using the subread-2.0.1 package (featureCounts) with a M31 GTF annotation file and the following parameters: “- s 2 -t exon -g gene_name”. Differential expression analysis was performed using DESeq2-1.28.1 with a P-value threshold of 0.05 within R version 4.0.2 (https://www.R-project.org/). A PCA plot was created within R using the pcaExplorer to measure sample expression variance. Heatmaps were generated using the pheatmap package were given the rLog transformed expression values. Gene ontology analyses were performed using the EnrichR package.

### Cell adhesion assay (Fig-7H)

Collagen-coated 96-well plates were prepared by diluting mouse collagen IV (R&D Systems, 3410-010-01 in sterile, cell culture-grade water to a concentration of 10μg/mL. 35μL of collagen was added to each well of a 96-well plate and allowed to dry in the hood under UV light for 2 hours. Each well was washed twice with wash buffer, (0.1% BSA in DPBS), blocked with 100μL 2% BSA in DPBS for 1 hour, followed by two more washes. A plate was prepared for each of the two timepoints: 30 and 60 minutes, respectively. 1 x 10⁶ HGS-3 cells were seeded in a 10cm dish for each concentration tested in complete DMEM (Gibco, cat#: 11885-084) supplemented with 10% FBS (Avantor# 89510-186), 1% penicillin/streptomycin (Gibco#: 15140-122). After allowing the cells to adhere overnight, each respective dish was treated with UR241-2 or DMSO vehicle (Vx) and incubated for 24 hours. Treated cells were harvested and washed twice in serum-free media before being counted and seeded into collagen-coated plates at 2000 cells/well in 100μL serum-free media. At each timepoint, each well was gently washed with wash buffer to remove non-adherent cells, leaving 100μL clean wash buffer in each well. Cells fixed by adding 100μL 10% TCA to each well without discarding existing supernatant and placed at 4°C for one hour before proceeding with SRB protocol.

### Cell migration/wound-healing assay (Fig-7I)

HGS-3 cell line was seeded in a 24-well plate, with 10,000 cells per well in complete DMEM (Gibco, cat#: 11885-084) supplemented with 10% FBS (Avantor# 89510-186), 1% penicillin/streptomycin (Gibco#: 15140-122). Once cells approached ∼70% cell density, they were treated with UR241-2 for 24 hours. After treatment, each well received a single vertical scratch using a P1000 tip. Scratched wells were rinsed once with media to remove detached cells and given fresh media. Initial images were taken immediately following this, represented by the “0” timepoint. Images were taken of the scratch at the same location in 12-hour intervals for 24 hours. Images were taken on an Olympus CKX41 inverted phase-contrast microscope with an Olympus DP74 camera and CellSens software. Image analysis was completed using Fiji’s ImageJ software and the wound healing size tool plugin developed by Suarez-Arnedo et al^52^.

### shRNA knockdown of IRAK4 in HGS-3 murine EOC cells

HGS-3 cells were transfected with murine shRNA plasmid (Santa Cruz Biotechnology, cat# sc-45401-SH) or the control plasmid (control shRNA Plasmid-A, cat#sc-108060) each using lipofectamine (Invitrogen, cat# L3000008) The media was changed after 1 day and cells were treated with increasing doses of puromycin antibiotic (2, 4, 6, 8 and finally 10µM). The surviving cells were collected and a portion of them were frozen in liquid nitrogen.

The remaining cells of both control plasmid and IRAK4^KD^ plasmid groups were lysed and expression of IRAK4 (IRAK4 antibody: cat#4363, dilution 1:1000, Cell Signaling Technology) was examined by western blot following the methods previously described. The same stripped membrane was probed with GAPDH (Cell Signaling Tech, cat #2118, dilution 1:10,000). The membranes were stripped again and probed with E-cadherin antibody (Cell Signaling Tech, cat #3195, dilution 1:500).

## Statistical analysis

Statistical significance between independent groups for non-repeated measurements was evaluated using the Wilcoxon-Mann-Whitney test, a nonparametric T-test. This method applies to outcomes measured on a continuous scale when small samples may hamper confirmation of distributional assumptions. Xenograft model data was analyzed using a repeated measures analysis of variance. The statistical model was fit using maximum likelihood estimation with treatment, day, and the interaction between treatment and day as fixed effects. Mouse was considered a random effect, and the correlation of repeated measures on the same mouse over time was handled using a first-order ante-dependence variance-covariance structure which varied by treatment. Model assumptions were verified graphically. If the Type 3 F-test for the interaction between treatment and day was significant, then treatment comparisons at each day were conducted using a T-test. Analysis was conducted using SAS v9.4 Proc Mixed (Cary, NC).

## Discussion

This study investigated the roles of IL-1β/IL1R1/IRAK4 in EOC seeding and tumor growth, specifically under the inflammatory signaling axis. We also show the therapeutic benefits of UR241-2, an IRAK4 inhibitor, in preventing and mitigating tumor growth at the site of needle injury or inflamed site in peritoneum. EOC metastasis, particularly peritoneal seeding, remains one of its most lethal aspects, especially during recurrence. Following the removal of the omentum in the initial surgery and subsequent chemotherapy, the peritoneum remains the primary site for metastatic and chemoresistant disease. Previously, the mechanism underlying EOC peritoneal seeding^53–56^ were partially understood, and models recapitulating these events were unavailable, and as a result, no targeted therapies could be developed to specifically target this lethal process. Now, our needle-injury induced MiM model, fills the need for a convenient, reproducible, non-surgical syngeneic animal-based surrogate model of cell adhesion and changes in ECM reorganization and immune milieu. The MiM model can facilitate mechanistic studies on ECM dynamics, immune environment interactions on a temporal basis, and can be used to screen compound libraries to select new therapeutics that can control EOC seeding. The MiM model is superior to that described by Jia et al^1^ which demonstrated EOC seeding on injured mesothelium and wounded ovaries but required surgical procedures on the mice.

Based on the roles of IL-1β/IL-1R1 and IRAK4 in EOC seeding at inflamed or injured sites, it is rational to use Anakinra (recombinant IL-1R1 antagonist), Canakinumab (anti-human IL-1β monoclonal antibody), or UR241-2 after initial treatment with surgery and chemotherapy to prevent, delay, or reduce metastatic burden upon recurrence. However, the lack of anti-tumor immune changes in tumors derived from IL-1R1 knockout) (KO) mice (Supplemental Fig-5 suggests that Anakinra or Canakinumab may not offer lasting responses. On the other hand, UR241-2 treatment modulated immune cell populations, particularly in the peritoneal cavity, an environment where immune cells can circulate and enter peritoneal metastases^57^. We noted an increase in MHCII-expressing macrophages and neutrophils, and a decrease in M2-like macrophages, which may contribute to a shift from an anti-inflammatory to a pro-inflammatory environment that may prevent tumor growth at the site of injury. This observation highlights the importance of immune modulation in cancer therapy, a strategy that has gained considerable attention in the field of immuno-oncology. In addition, UR241-2 has demonstrated notable pharmacologic and ADMET attributes, including inhibition of IL-1β-induced IRAK4 phosphorylation, reduction of the activation of long-form IRAK4 without affecting its short-form isoform, revealing various nuanced but related mechanisms of actions, inhibition of NF-κβ signaling, a key downstream pathway of IL1/TLR/IRAK4 signaling, reduction of the adhesion, proliferation, and division of EOC cells, with significant effects on colony formation, and reduction of the growth of SKOV-3 xenograft and syngeneic tumors, without adverse effects on body weight or hematology. Further, as an indicator of excellent pharmacokinetics in mice, IP administered 20mg/kg UR241-2 maintains ∼8pM of inhibitor in plasma for 8 hours which indicates that drug is stable in C57Bl/6 mice and drug responses seen in both xenograft and syngeneic animal models correlates with plasma levels (Supplementary Figure-3G).

The clinical scopes of both UR241-2/other IRAK4 inhibitors and Anakinra and Canakinumab lies primarily in combination with existing chemotherapies and/or immunotherapies, that too preferably after first round of chemotherapy and surgery. Increased sensitivity of KRAS-mutant colon cancer to fluorouracil treatment when combined with Anakinra is an example^58^. Anakinra was additionally tested as the AGAP (Gemcitabine, nab-paclitaxel, cisplatin, and anakinra) regimen in patients with localized pancreatic ductal adenocarcinoma (PDAC)^59^. Similarly, CA4948, an IRAK4 inhibitor is being tested in combination with PD-L1/PD-1 in pancreatic, esophageal/gastric, bladder, AML and other malignancies. For EOC treatment, our future studies will examine the combination of UR241-2+Paclitaxel. Paclitaxel activates the TLR4 pathway which promotes chemoresistance. The combination of paclitaxel+IRAK4i UR241-2 will likely block TLR4 orchestrated resistance signaling and likely increase the response of paclitaxel improving the overall survival.

Taken together, while this study provides much-needed insights behind cell adhesion at the injured mesothelium, genomic changes and the resultant tumor growth, and provides a therapeutic candidate UR241-2 to control it, further research is needed in greater details to fully elucidate the complexities of EOC adhesion, inflammation, and the tumor immune-milieu at play, and to advance UR241-2 to the EOC clinics. Notably, our study does not reveal the role of IL1β/IL1R1/IRAK4 in EOC cell’s attachment on the omentum. Since omentum resident macrophages promote the migration and colonization of EOC cells^60^, we could not identify if UR241-2 treatment interfered with the polarization of omental macrophages into a pro-inflammatory or anti-inflammatory phenotype, which would be important in identifying macrophage changes within the tumor itself. Since Il1/Il1r1^LoF^ or UR241-2 treatment did not reduce the tumor burden on omentum in our animal models, we infer that omental adhesion of EOC cells may occur independent of IL-1β/IL-1R1/IRAK4 axis. However, since surgical removal of the omentum is commonly performed in EOC treatment, the peritoneum, not the omentum, remains the clinically relevant site to monitor and control the future emergence of EOC.

## Supporting information

revised supplemental

## Acknowledgement

Empire Discovery Institute (EDI) paid for Pan-kinase screening and ADMET studies. EDI played no roles in writing of this manuscript. ChatGPT 4o was used in editing the sentences written by the authors to improve readability. ChatGPT suggested changes in the sentences were verified for scientific accuracies. Dr Christina Annunuziata MD PhD provided OVCAR-3_NF-kβ cell-line.

## Authors contributions

JPM performed flow cytometry and edited the manuscript. NK performed western blotting. CWS performed cell adhesion and migration assays, microscopy, and analyzed/quantified images. NAS performed docking and simulations. EL performed animal experiments and did the treatments, measured tumor sizes and animal weights and performed CBC analysis repeatedly. MS performed the statistical analysis. KKK helped develop IRAK4^KD^ HGS-3 cell clones and performed NF-kβ reporter assay. MA and RK designed and partially funded ADME assays. SS analyzed the flow data under supervision of JB. CMA provided OVACAR-3-NF-κB reporter cell-line. JA, CB and EP performed bulk-seq. MKK directed the synthesis of UR241-2 and analogs. RRT, RGM provided laboratory resources, funding and critically read and edited the manuscript multiple rounds. MWB provided KO animals, guidance and reviewed the data. RKS brought resources with RGM and MWB, conceived the study, designed and synthesized UR241-2 first round and wrote the manuscript.

