## Supplementary material for "IL-1β/IRAK4 Axis Promotes Ovarian Tumor Development at the Mesothelium Injury Sites": revised supplemental

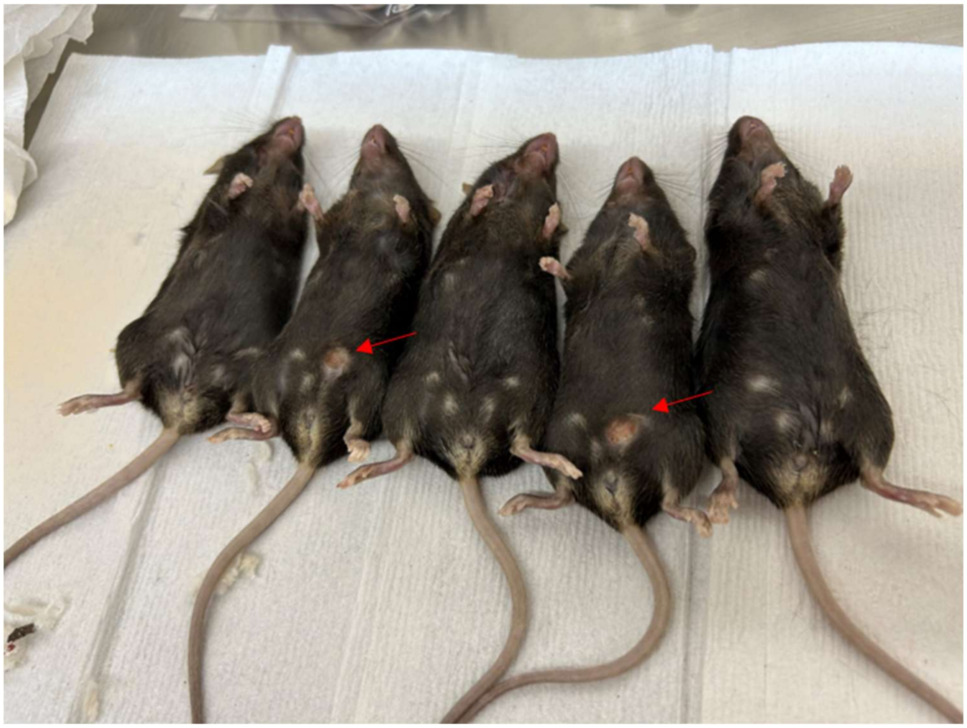

**Supplementary Figure-1:** HGS-3 murine high-grade serous EOC cells (3 million/per mice) were implanted intraperitoneally using 21-gauge needle in C57BL/6 WT and C57BL/6 NLRP3$^{KO}$ mice. Mice were observed for 45-50 days and euthanized. Tumors formed on needle injury site, protruding at the skin as well as in the peritoneum and on the omentum, shown by red arrows, were isolated, weighed and frozen in liquid -nitrogen. Lavages via washing with sterile PBS(5mL) were also collected. The studies were repeated twice. A representative experiment is shown. Weights of the omental did not differ between C57BL/6 WT and NLRP3$^{KO}$ mice. Similarly, the tumor sizes at the site of injury did not differ between C57BL/6 WT and NLRP3$^{KO}$ mice.

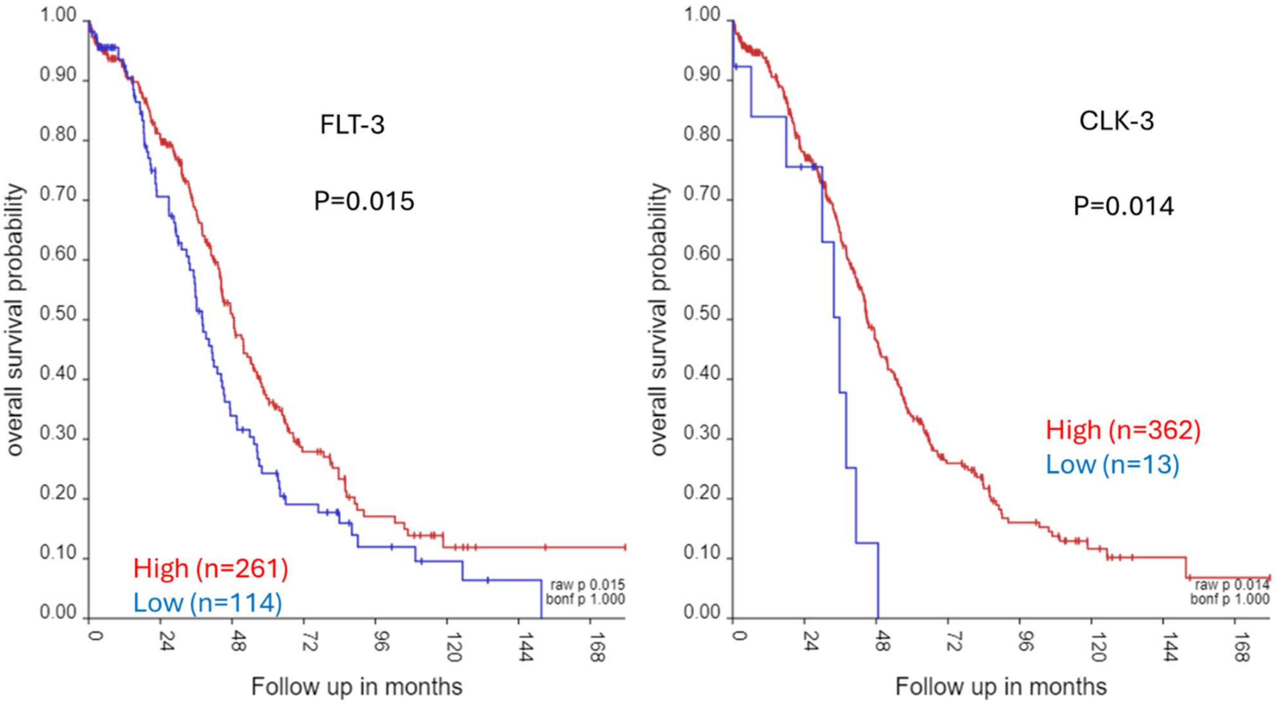


**Supplementary Figure-2:** Analysis of ovarian serous cystadenocarcinoma (2022-v32) microarray data (TCGA-381-tpm-gencode36) of ovarian cancer patients using R2 Genomics Analysis and Visualization Platform tools showed that FLT-3 and CLK-3 mRNA overexpression predicts poor survival.

A

| Compound ID | λ (nm) | Solubility (μM) |
|---|---|---|
| UR-241-2 | 280 | 29 |
| Verapamil | 280 | 89 |
| Tamoxifen | 280 | 4.3 |

B

| Project Compound ID | Human Liver Microsomes | | | Mouse Liver Microsomes | | |
|---|---|---|---|---|---|---|
| | $t_{1/2}$ (min) | $CL_{int}$ (μL/min/mg) | $E_H$ | $t_{1/2}$ (min) | $CL_{int}$ (μL/min/mg) | $E_H$ |
| EDI-238382 | 214 | 3.24 | 11% | 13.5 | 51.4 | 53% |
| Testosterone* | 21.8 | 31.8 | 56% | 7.59 | 365 | 89% |
| UR241-2 | 208 | 3.3 | 12% | 8.70 | 79.7 | 63% |
| Testosterone* | 11.3 | 61.5 | 71% | 6.83 | 406 | 90% |

C

| Compound ID | CYP $IC_{50}$ (μM) | | | | | | | |
|---|---|---|---|---|---|---|---|---|
| | CYP1A2 | CYP2B6 | CYP2C8 | CYP2C9 | CYP2C19 | CYP2D6 | CYP3A4M | CYP3A4T |
| UR-241-2 | >100 | >100 | 79 | 30 | 45 | >100 | 58 | 39 |
| Furafylline | 1.8 | | | | | | | |
| Ticlopidine Hydrochloride | | 0.093 | | | | | | |
| Montelukast Sodium Hydrate | | | 0.013 | | | | | |
| Sulfaphenazole | | | | 0.17 | | | | |
| (+)-N-3-Benzylnirvanol | | | | | 0.20 | | | |
| Quinidine | | | | | | 0.11 | | |
| Ketoconazole | | | | | | | 0.014 | 0.010 |
| Z' | 0.73 | 0.80 | 0.78 | 0.77 | 0.74 | 0.66 | 0.79 | 0.76 |
| $r^2$ | 1.0 | 0.99 | 0.99 | 0.99 | 1.0 | 0.99 | 1.0 | 1.0 |

| % control activity at 100 μM | | | | | | | |
|---|---|---|---|---|---|---|---|
| CYP1A2 | CYP2B6 | CYP2C8 | CYP2C9 | CYP2C19 | CYP2D6 | CYP3A4M | CYP3A4T |
| 68 | 67 | 42 | 28 | 33 | 62 | 49 | 40 |

D

| Project Compound ID | Protein Binding, % | | Recovery, % | |
|---|---|---|---|---|
| | Human Plasma | Mouse Plasma | Human Plasma | Mouse Plasma |
| *UR-241-2 | 97.4 ± 0.3 | 93.8 ± 0.8 | 92.1 | 55.1 |
| Propranolol | 73.7 ± 1.6 | 81.2 ± 1.3 | 100 | 115 |

E

| Compound ID | Lot | % Recovery | | $P_{app}$ ($x10^{-6}$ cm/s) | | Efflux Ratio | Permeability Classification | Significant Efflux |
|---|---|---|---|---|---|---|---|---|
| | | A-B | B-A | A-B | B-A | | | |
| UR-241-2 | 12-WEN-111-1 | 76.8 | 85.2 | 11.9 | 49.4 | 4.14 | High | Yes |

F

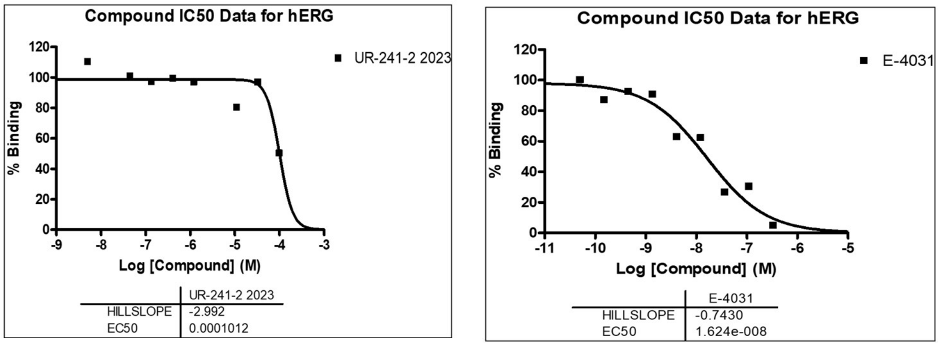


G

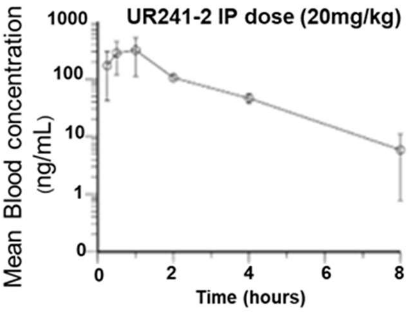

**Supplementary Figure-3**: (**A**): kinetic solubility of UR241-1 is shown. (**B**): Quantified stability of UR241-2 in human and mouse liver microsomes is shown. (**C**): Inhibition of CYP450 isoforms by UR241-2 is quantified using HPLC. Controls used were furafyllin, Ticlopidine HCL, Montelukast sodium hydrate, sulfaphenazole, N-3-benznirvanol, Qunidine, ketoconazole. CYP isoforms affected by UR241-2 in terms of %-inhibition are shown. (**D**): Human and murine plasma protein binding of UR241-1 is shown in % units. Propranolol was used as a control. (**E**): CaCo-2 cell permeability of UR241-2 is shown. Efflux ratio of 4.14 indicates that UR241-2 faces significant efflux. (**F**): UR241-2 does not inhibit hERG. $IC_{50}$ is 101.2µM. E-4031 was used as control. $IC_{50}$ for E-4031 was $1.62e^{-08}$. (**G**): PK of UR241-2 at 20mg/kg administered IP is shown. It is shown that ~8ng/ml concentration of UR241-2 is maintained up until $8^{th}$ hour of testing.

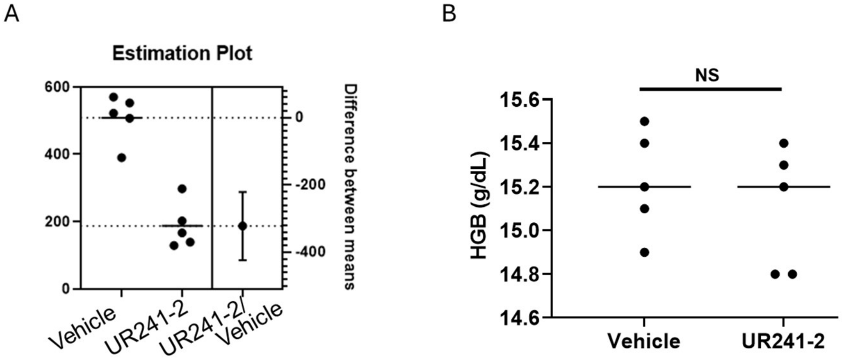


**Supplementary Figure-4**: (**A**): The statistical difference between the vehicle and UR241-2 treated tumors was also analyzed by estimation plot using GraphPrism version 8.0 is shown. (**B)**: Analysis of the peripheral blood showed that hemoglobin (HB) levels did not differ between the vehicle and UR241-2 treated mice.

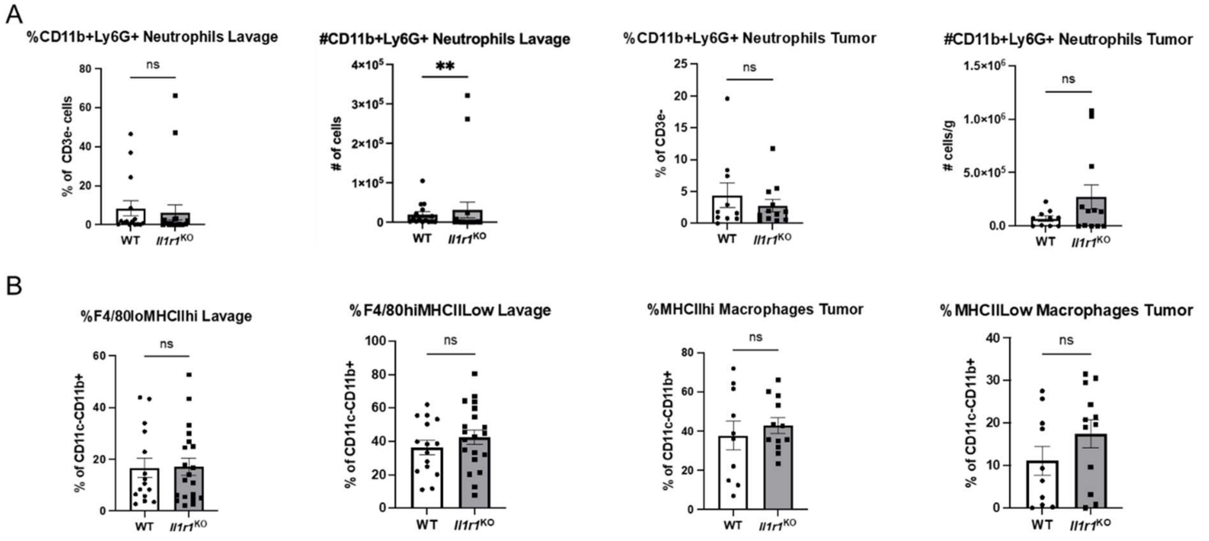


**Supplementary Figure-5:** Il1r1 KO does not significantly impact myeloid percentages in the peritoneal lavage and tumor. A) Peritoneal lavage and tumor percentages and cell numbers of CD11b+Ly6G+ neutrophils from flow cytometry data of wild type and IL1R1 KO mice injected with HGS3 tumors. B) Peritoneal lavage percentages and cell numbers of F4/80loMHCIIhi and F4/80hiMHCIILow macrophages from the peritoneal lavage and percentage of MHCIIhi and MHCIILow macrophages from tumors from flow cytometry data of wild type and IL1R1 KO mice injected with HGS3 tumors. N = 15-20 mice per group from 3 independent experiments. Statistical significance was determined using Mann-Whitney test, *p < 0.033, **p < 0.002, ***p < 0.001

| Reference /Cat | Manufacturer | Name | Other Name | Clone | Fluorochrome | Max Excitation | Max Emission | Laser |
|---|---|---|---|---|---|---|---|---|
| 565992 | BD Biosciences | CD3e | | 145-2C11 | **BUV395** | **348** | **395** | **UV** |
| 364-0081-82 | eBioscience | CD8a | | 53-6.7 | **BUV496** | | | **UV** |
| 612793 | BD Biosciences | CD69 | | H1.2F3 | **BUV737** | **350** | **737** | **UV** |
| 749284 | BD OptiBuild | F4/80 | | T45-2342 | **BUV563** | **351** | **561** | **UV** |
| 750042 | BD OptiBuild | CD192 | CCR2 | 475301 | **BUV661** | **348** | **661** | **UV** |
| 568287 | BD Horizon | CD19 | | 1D3 | **BUV805** | | | **UV** |
| 752299 | BD OptiBuild | CD279 | PD-1 | J43 | **BUV615** | **350** | **616** | **UV** |
| 563053 | BD Horizon | CD45 | | 30-F11 | **Brilliant Violet 605** | **407** | **605** | **Violet** |
| 564023 | BD Biosciences | CD25 | | PC61 | **Brilliant Violet 785** | **408** | **786** | Violet |
| 101251 | Biolegend | CD11b | | M1/70 | **Brilliant Violet 421** | **408** | **422** | **Violet** |
| 560458 | BD Horizon | CD11b | Gr1 | 1A8 | **V450** | **404** | **448** | **Violet** |
| 560603 | BD Horizon | Ly6g | Gr1 | 1A8 | **V450** | **404** | **448** | **Violet** |
| 560593 | BD Pharmingin | Ly-6C | | AL-21 | **PE-Cy7** | **568** | **778** | **Yellow-Green** |
| 558091 | BD Biosciences | CD274 | PD-L1 | MIH5 | **PE** | **496** | **578** | **Yellow-Green** |
| 11-5773-82 | eBioscience | FoxP3 | | FJK-16s | **FITC** | **494** | **520** | **Blue** |
| 566504 | BD Horizon | CD11c | | HL3 | **BB700** | | | **Blue** |
| | Biolegend | CX3CR1 | Fractalkine receptor | SA011F11 | **PE-Dazzle594** | | | **Blue** |
| 552051 | BD Pharmingin | CD4 | | GK1.5 | **APC-Cy7** | **650** | **775** | **Red** |
| 17-5321-82 | eBioscience | MHCII | I-A/I-E | M5/114.15.2 | **APC** | **650** | **660** | **Red** |
| 47-5932-82 | invitrogen | Ly6C | | HK1.4 | **APC-eFluor 780** | | | |
| 25-2061-82 | invitrogen | CD206 | | MR6F3 | **PE-Cy7** | **568** | **778** | **Yellow-Green** |
| L34959 | Invitrogen | LiveDead Yellow | | | | | | **Violet** |

**Supplementary data-6:** List and details of flow cytometry antibodies used in this study

**Supplementary Data-7**

**NMR and Mass spectrometry data for UR241-2:**

$^{1}H$ NMR (400 MHz, $CDCl_3$), δ: 10.52 (s, 1H), 8.51 (d, *J* = 8.7 Hz, 1H), 8.21 (d, *J* = 6.6 Hz, 1H), 8.04 – 7.93 (m, 2H), 7.72 (d, *J* = 2.1 Hz, 1H), 6.98 (d, *J* = 2.1 Hz, 1H), 6.60 (dd, *J* = 8.8, 2.5 Hz, 1H), 6.56 (d, *J* = 2.4 Hz, 1H), 4.01 (s, 3H), 3.86 – 3.77 (m, 4H), 3.21 – 3.11 (m, 4H). LCMS (m/z): 428.3 $[M+H]^+$.

**NMR and Mass spectrometry data for PSP-0099:**

$^{1}H$ NMR (400 MHz, $CDCl_3$), δ: 10.51 (s, 1H), 8.48 (d, J = 9.3 Hz, 1H), 8.21 (d, J = 7.2 Hz, 1H), 8.02 – 7.92 (m, 2H), 7.72 (d, J = 1.9 Hz, 1H), 6.98 (s, 1H), 6.60 (d, J = 6.5 Hz, 2H), 4.00 (s, 3H), 3.57 – 3.50 (m, 3H), 2.86 – 2.77 (m, 4H). LCMS (m/z): 396.3 $[M+H]^+$.

**NMR and Mass spectrometry data for PSP-0100:**

$^{1}H$ NMR (400 MHz, $CDCl_3$), δ: 10.51 (s, 1H), 8.49 (d, J = 8.7 Hz, 1H), 8.21 (d, J = 7.3 Hz, 1H), 7.97 (dt, J = 15.3, 7.5 Hz, 2H), 7.72 (d, J = 1.5 Hz, 1H), 6.98 (s, 1H), 6.64 (dd, J = 8.8, 2.4 Hz, 1H), 6.60 (d, J = 2.4 Hz, 1H), 3.99 (d, J = 10.2 Hz, 3H), 3.95 (dd, J = 10.2, 3.2 Hz, 2H), 3.54 (dd, J = 18.1, 3.7 Hz, 2H), 3.00 – 2.85 (m, 4H). LCMS (m/z): 412.2 $[M+H]^+$.
